# Peptide signaling in the paraventricular thalamus contributes to disrupted adult reward behaviors after early-life adversity

**DOI:** 10.64898/2026.03.31.715695

**Authors:** Amalia Floriou-Servou, Ryan Weber, Yuncai Chen, Cassandra L. Kooiker, Madison R. Tetzlaff, Matthew T. Birnie, Annabel K. Short, Roshae Roberts, Heidi Y. Liang, Magdalena Gantuz, Mason Hardy, Ali Mortazavi, Tallie Z. Baram

## Abstract

Early-life adversity (ELA) is strongly linked to emotional disorders characterized by dysregulated reward behaviors including diminished motivation for and enjoyment of rewards (anhedonia). However, how <u>transient</u> ELA leads to enduring disruptions of reward behaviors remains unclear. Using activity-dependent genetic labeling during ELA, we previously found that the paraventricular thalamic nucleus (PVT) uniquely distinguishes ELA from typical rearing, and inhibiting ELA-activated PVT cells in adult ELA mice rescued reward behaviors in a sex-specific manner. However, the cellular mechanisms by which ELA alters the function of PVT neurons enduringly, to impact adult reward behaviors is unresolved. Here, we probed potential gene expression mechanisms by assessing reward-induced gene translation selectively in early-life activated (TRAPed) PVT cells, and identified genes regulated by reward in a sex-specific way. Combining data- and hypothesis-driven approaches, we targeted the corticotropin-releasing hormone receptor type 1 (CRHR1) as potentially mediating ELA-induced disruptions of reward behaviors. We find that CRISPR-Cas9-mediated *Crhr1* deletion in TRAPed-PVT cells restores typical reward behaviors in ELA mice. Our results provide insight into the developmental origin of mental illness and identify potentially powerful molecular targets for prevention or mitigation of the consequences of ELA on reward behaviors germane to human mental illness.

## Main

Early life adversity (ELA) affects a majority of the world’s children (Madigan et al., 2025; Merrick et al., 2018) and is associated with negative emotional outcomes and elevated risk for mental illness later in life (Birnie & Baram, 2025; Fleming et al., 2025; McKay et al., 2022; Nelson et al., 2025). Specifically, ELA has been linked to disorders such as depression and substance use, which are characterized by dysregulation of the reward system (Gaffrey et al., 2018; Kopala-Sibley et al., 2018; Novick et al., 2018). In animal models, ELA directly causes changes in the reward circuitry (Birnie et al., 2020; Bolton, Molet, et al., 2018), in reward responsiveness (Kangas et al., 2021) and in reward behaviors, and the effects are sex-dependent. Males raised in ELA show reduced interest in pleasure (anhedonia) (Birnie et al., 2023; Bolton, Ruiz, et al., 2018; Hall et al., 2025; Levis et al., 2022; Phillips, 2024; Scheggi et al., 2018), while ELA females have increased motivation for both natural and drug rewards (Birnie et al., 2023; Kooiker et al., 2024; Levis et al., 2021), compared to animals reared in typical conditions.

How does transient early life adversity lead to long-term alterations in brain function and behavior? Understanding the underlying mechanisms is required for preventing or treating mental illnesses related to ELA. ELA may impact brain function enduringly through long-term changes in gene expression. Indeed, several studies point out to epigenetic and transcriptional changes induced by ELA in several stress-sensitive brain regions including the hypothalamus (Short et al., 2023), ventral tegmental area (Peña et al., 2017) and hippocampus (Bolton et al., 2020; Kos et al., 2023; Malave et al., 2022). However, the specific brain regions and cell types or subtypes that initially sense and encode ELA are unknown (Weber et al., 2025), especially when ELA is experienced during the first weeks of life when hippocampal memory systems are not yet operational (Bisaz et al., 2014; Travaglia et al., 2016). Epigenomic changes in these regions may underlie disrupted adult reward behaviors.

Using activity-dependent genetic labeling during the ELA period, we found robust neuronal activation in the paraventricular thalamic nucleus (PVT) during the first week of life. Notably, PVT was the only brain area among the ones tested to distinguish ELA from typical rearing (Kooiker et al., 2023). The PVT is a key node bridging cortical and subcortical networks to assess reward and threat cues within the context of prior stress (Bhatnagar & Dallman, 1998; Dong et al., 2017; Kooiker et al., 2021; Li & Kirouac, 2008; McGinty & Otis, 2020). The region is thus posed to both encode ELA (Do-Monte et al., 2015; Heydendael et al., 2011) and influence its function throughout life, including behaviors in response to reward cues.

Here, we test the hypothesis that ELA activates a subset of PVT neurons, leading to enduring changes in gene expression which, in turn, govern the dysregulated reward behaviors in adult ELA mice. Leveraging genetic tagging (Targeted Recombination in Active Populations, TRAP2, DeNardo et al., 2019), combined with ribosome trapping (Translating Ribosome Affinity Purification, TRAP, Heiman et al., 2014), we determined actively translated mRNA in response to reward cues in cells activated during ELA or control rearing (TRAPed cells). Reward-induced gene translation in TRAPed neurons was significantly modified by rearing and further modulated by sex, in line with the behavioral phenotypes. Combining hypothesis-and data-driven approaches, and building on enriched TRAPing of corticotropin-releasing hormone (CRH) receptor-expressing cells during ELA, we chose to focus on the stress- and reward-reactive CRH signalling via this receptor as a target (Birnie et al., 2023). Indeed, the use of CRISPR-mediated deletion of CRHR1 exclusively in TRAPed-PVT cells, had no effect on baseline behaviors, and yet normalized reward behaviors in both male and female adult ELA mice.

## Results

### ELA changes adult motivated behaviors in a sex specific way, and directly engages the PVT

To determine the impact of ELA on reward behaviors, we exposed adult mice of both sexes that were reared under typical conditions (controls; CTL) or experienced one week of ELA, to palatable food (cocoa pebbles; 1h per day; 4 consecutive days), and measured consumption. The ELA paradigm used here involves chronic stress of pups and dams accomplished by limiting access to bedding and nesting in home cages between postnatal days 2-10 (P2-P10) (Molet et al., 2016; Rice et al., 2008; Birnie et al., 2023). Adult male ELA mice consumed less palatable food compared to CTL (Fig. 1A-B). By contrast, ELA females consumed more palatable food per body weight compared to their CTL counterparts (Fig. 1C-D and S1A). In these non-food restricted mice, consumption of laboratory chow did not differ by rearing (Fig. S1B-E) indicating that ELA influenced hedonic rather than nutritional feeding behaviors.

**Fig. 1.**
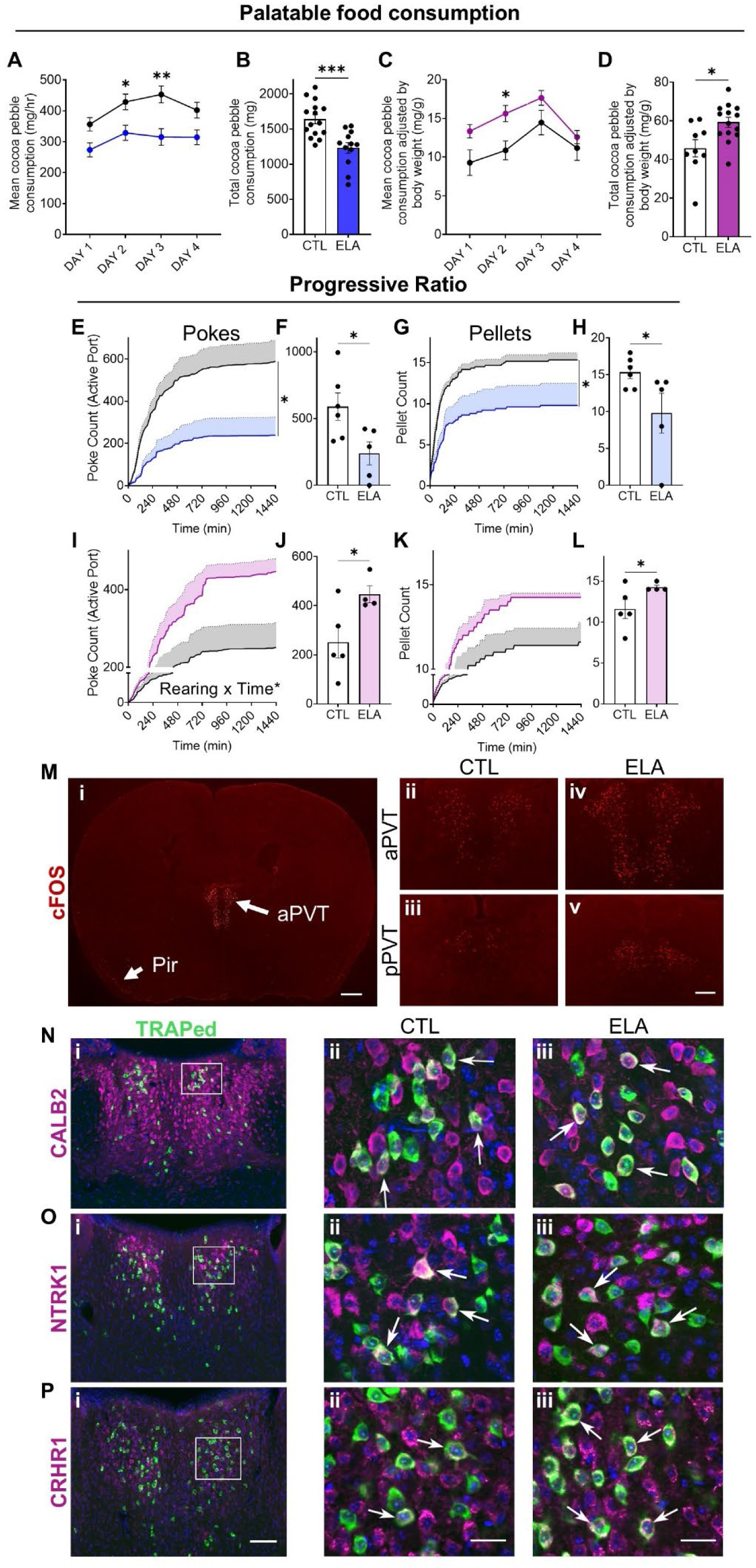
ELA changes adult motivated behaviors in a sex-specific way, and directly engages the PVT. **A-B.** Male mice with prior ELA experience consumed less palatable food compared to CTL mice (cocoa pebble weight consumed/hour over 4 days, main effect of Rearing: F(1,25) = 16.81, P = 0.0004, two-way ANOVA with Sidak post hoc tests (**A**) and total consumption: t(25) = 4.099, P = 0.0004, unpaired t test (**B**)). N = 15 CTL & 12 ELA males **C-D.** Female ELA mice consumed more palatable food compared to CTL (cocoa pebble weight adjusted by body weight, consumed/hour over 4 days, main effect of Rearing: F(1,21) = 7.765, P = 0.0111, two-way ANOVA with Sidak post hoc tests (**C**) and total consumption adjusted by body weight: t(21) = 2.792, P = 0.0109, unpaired t test (**D**)). Consumption was adjusted by body weight because ELA females weighed singificantly less than CTL (Fig. S1). N = 9 CTL & 14 ELA females **E-H.** ELA males poked the active port fewer times and received fewer sugar pellets compared to CTL, on a progressive ratio test with FED3 devices (nose pokes over 24 hours, main effect of Rearing: F(1,9) = 7.268, P = 0.0246, two-way ANOVA (**E**) and total nose pokes: t(9) = 2.553, P = 0.0311, unpaired t test (**F**); sugar pellets retrieved over 24 hours, main effect of Rearing: F(1,9) = 5.393, P = 0.0453, two-way ANOVA (**G**) and total pellets retrieved: t(9) = 2.127, P = 0.0312, one-tailed unpaired t test (**H**)). N = 6 CTL & 5 ELA males **I-L.** ELA females poked the active port more times as the effort requirement increased, and received more sugar pellets compared to CTL, on a progressive ratio test with FED3 devices (nose pokes over 24 hours, interaction Rearing*Time: F(1.898, 13.28) = 4.457, P = 0.0346, main effect of Rearing: F(1,7) = 4.447, P = 0.0729, two-way ANOVA (**I**) and total nose pokes: t(7) = 2.489, P = 0.0417, unpaired t test (**J**); sugar pellets retrieved over 24 hours, main effect of Rearing: F(1,7) = 4.556, P = 0.0702, two-way ANOVA (**K**) and total pellets retrieved: t(7) = 1.977, P = 0.0443, one-tailed unpaired t test (**L**)). N = 5 CTL & 6 ELA females **M.** cFOS expression in the brain of a postnatal day 10 (P10) mouse. Microscopy image of a coronal section (i) and higher magnification images of the anterior PVT and posterior PVT from a CTL (ii-iii) and ELA (iv-v) mouse. Scale bar in (i) = 500 μm and in (ii-v) = 125 μm **N-P.** Colocalization of TRAPed cells with CALB2 (**N**), NTRK1 (**O**) and CRHR1 (**P**) in the PVT of CTL (ii) and ELA (iii) mice. Scale bar in (i) = 100 μm and in (ii-iii) = 20 μm

To gain insight into the motivational and consummatory aspects of the hedonic feeding changes induced by ELA we tested mice under high-effort conditions, using the Feeding Experimentation Devices (FED3, Matikainen-Ankney et al., 2021) and a progressive ratio design. The FED3 dispenses sugar pellet rewards in response to nose-pokes (effort). Initially, a single pellet was dispensed after each nose-poke in the active port (fixed ratio [FR]1). Next, the device was set on a progressive ratio, requiring increasingly more pokes for the same reward. All groups learnt the task equally well preferring the active over the inactive port (Fig. S1F-K). In the progressive ratio task, male ELA mice poked less (Fig. 1E-F) and received fewer pellets compared to CTL (Fig. 1G-H). In contrast, ELA female mice poked more and received more pellets compared to CTL (Fig. 1I-L). These results implicate sex-specific motivational dysregulation of reward behaviors in adult ELA mice (Birnie et al., 2023; Kooiker et al., 2024; Levis et al., 2021).

We next sought to characterize neurons activated preferentially during ELA as measured by the expression of the immediate early gene *cFos*. *cFos*, in addition to signifying neuronal activation, is a major regulator of gene expression programs during development (Velazquez et al., 2015; West et al., 2002), and in response to stimuli (Greenberg & Ziff, 1984; Nestler et al., 2001; Phillips III et al., 2023). We first validated, using recapitulated immunohistochemistry, the ELA-dependent cFOS expression in the PVT established before using TRAP2 mice (Kooiker et al., 2023) (Fig. 1M). cFOS was robustly expressed throughout both the anterior and posterior PVT (Fig. 1Mii-iii), and much more weakly elsewhere (Fig. 1Mi). As expected, more cells expressed cFOS in ELA mice compared to CTL (Fig. 1Mii-v), in accord with our prior findings using activity-dependent genetic tagging (Kooiker et al., 2023). We then characterized the cell type of TRAPed PVT neurons, and found that, as expected. they expressed calretinin (CALB2; Fig. 1N), a hallmark of PVT neurons (Gao et al., 2023; Mátyás et al., 2018; Shima et al., 2023; Tetzlaff et al., 2025). Subsets of TRAPed neurons expressed neurotrophic tyrosine kinase receptor type 1 (NTRK1; Fig. 1O) and the CRH receptor type 1 (CRHR1; Fig. 1P). While the proportion of TRAPed cells expressing CALB2 and NTRK1 was similar in ELA and CTL PVT, the fraction of CRHR1-expressing neurons among all TRAPed cells was significantly larger in ELA mice compared to CTL (Fig. 1Pii-iii).

### Non-stimulated gene translation is minimally influenced by early-life rearing

Testing the hypothesis that PVT neurons differentially activated by ELA vs. typical rearing (TRAPed neurons) contribute to the dysregulated behaviors in response to reward cues in adult ELA mice requires access to genes actively translated in response to these cues. Therefore, we combined the TRAP2 mice with Translating Ribosome Affinity Purification (TRAP), by crossing the Fos-Cre^ERT2^ mice with those expressing the ribosomal tag EGFP-L10a in a Cre-dependent manner (nuTRAP mice). We induced CRE with tamoxifen administration during the ELA or CTL-rearing period at P6 to access cells activated during the subsequent 24-36 hours (Fig. 2A). We confirmed the expression of the EGFP tag in mouse brain in a pattern similar to the cFOS expression, and specifically a robust expression in the PVT (Fig. 2B-C). RNA isolated after immunopurification for the EGFP tag (post-IP) was significantly enriched for PVT genes (Fig. 2D-F) compared to pre-immunopurification (pre-IP). Specifically, differential gene expression analysis (DGEA) identified 374 upregulated genes (FDR<0.05, log2FC>1), and 2783 downregulated genes (FDR<0.05, log2FC< −1) in the post-IP fraction (Fig. 2E-F). Among the most strongly upregulated genes were those highly expressed in the PVT compared to neighboring thalamic nuclei, including *Calb2, Snca, Prph, Gal, Gck, Crhbp* and *Ntrk1* (thalamoseq database, (Phillips et al., 2019) (Fig. 2F and S2A). In contrast, genes primarily expressed in neighboring thalamic nuclei such as the mediodorsal nucleus (MD), the intermediodorsal nucleus (IMD) and the habenula (Hb) (Gao et al., 2023; Phillips et al., 2019) were downregulated post-IP (Fig. 2F and S2B). Post-IP fractions were enriched in neuronal markers (*Syn1* and *Syn2)* and depleted of markers of other cell types (Fig. S2C) indicating that most RNA was isolated from neurons. Finally, among PVT cell-type markers (Gao et al., 2023; Shima et al., 2023), genes encoding the CRH binding protein, *Crhbp* and *Htr1d* were enriched in TRAPed-PVT cells (Fig. S2D). GO term analysis showed that genes distinguishing the immunopurified TRAPed cell fraction are involved in catecholamine transmission, response to cocaine, feeding behavior and maternal behavior (Fig. S2E), all established PVT functions (Dumont et al., 2022; James et al., 2010; Watarai et al., 2020; Ye et al., 2022).

**Fig. 2.**
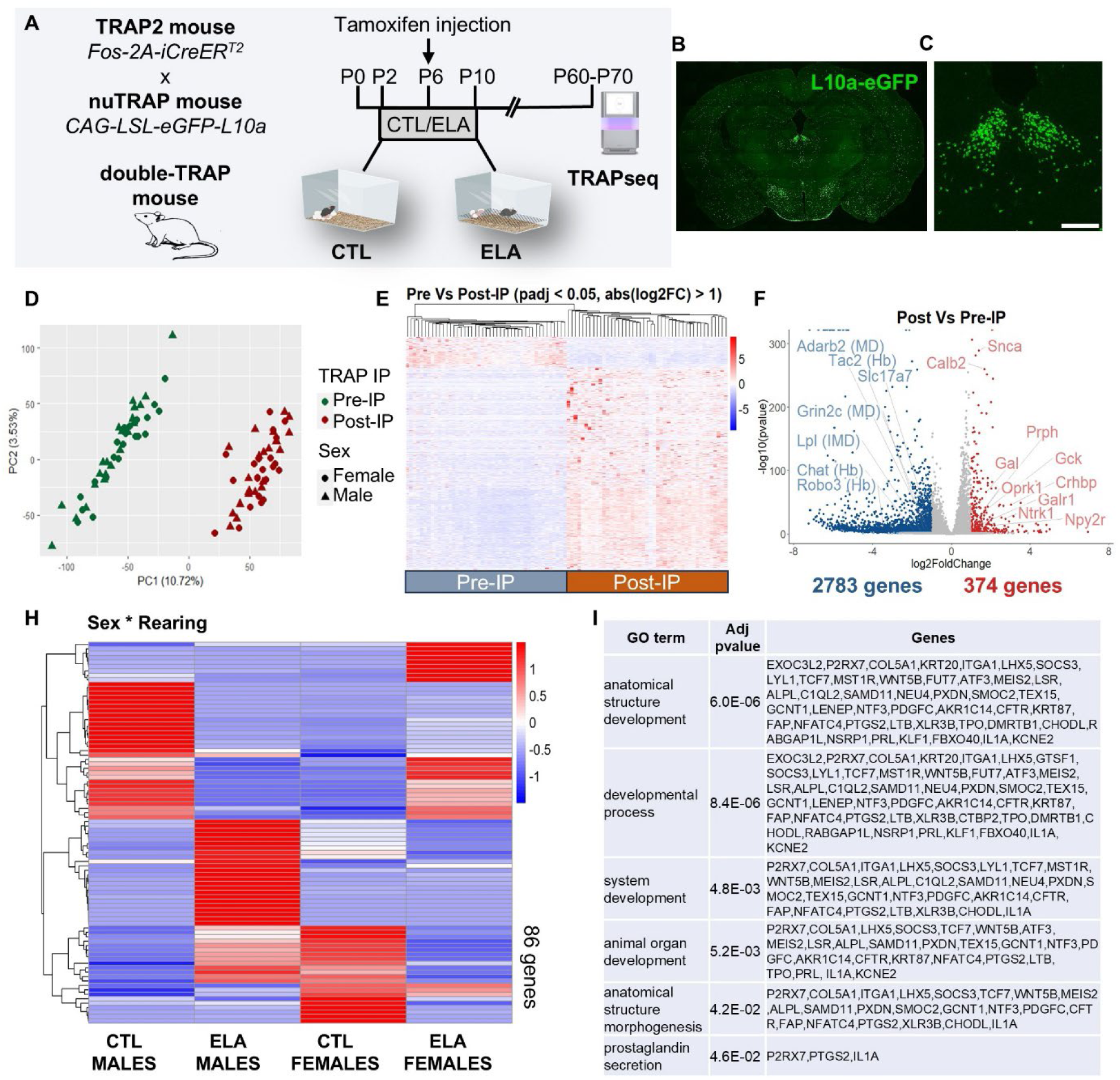
Non-stimulated gene translation is minimally influenced by early-life rearing. **A.** Experimental design for activity-dependent genetic labeling of cells activated in the first week of life, leading to permanent expression of a ribosomal eGFP tag, used for translating ribosome affinity purification (IP) and sequencing of the bound RNA (TRAPseq) in adulthood. CRE is induced by tamoxifen administration at P6, while mice are raised in CTL or ELA cages. **B-C.** Representative microscopy images showing that the PVT is effectively labeled with the ribosomal tag eGFP-L10a in double-TRAP mice (age P30). Coronal brain section (**B**) and higher magnification showing the PVT (posterior) (**C**). Scale bar = 200 μm **D.** Principal component analysis showing the variance between the pre- and post-IP fractions of 45 samples from 23 male and 22 female mice, following TRAPseq. **E-F.** Differentially expressed genes (DEGs) between the pre- and post-IP fractions following TRAPseq. Heatmap showing the expression levels of 3,157 differentially expressed genes (DEGs) in all samples (**E**). Volcano plot showing DEGs in the post-IP fractions compared to pre-IP (**F**). PVT-specific genes such as *Calb2*, *Snca*, Crhbp, *Galr1*, *Ntrk1*, and *Npy2r* have higher expression post-IP, while genes specific for neighbouring nuclei such as *Adarb2* (highly expressed in mediodorsal nucleus (MD)), *Lpl* (intermediodorsal nucleus (IMD)), *Slc17a7* (most thalamic nuclei except PVT) and *Tac2* (habenula (Hb)) have lower expression. Upregulated genes shown in dark red: FDR<0.05, log2FC>1, downregulated genes showed in dark blue FDR<0.05, log2FC< −1. **H.** Heatmap showing the averaged expression of the DEGs for the interaction Sex*Rearing (FDR<0.05, log2FC>0.5 or log2FC< −0.5). N = 10 CTL & 13 ELA males, 12 CTL & 10 ELA females **I.** GO term analysis of the 86 DEGs for the interaction Sex*Rearing.

We next assessed differences in actively translated genes between ELA and CTL mice of both sexes. While ELA dramatically modified behaviors in response to reward cues, typical daily behaviors of adult ELA mice were not obviously changed. Accordingly, there were limited rearing-dependent differences in actively translated genes in TRAPed-PVT cells from mice that were sacrificed at rest. Differential gene expression (DEG) analysis identified 34 upregulated and 25 downregulated genes in males (FDR<0.05 and log2FC>0.5 or log2FC< −0.5, Fig. S3A), and 18 upregulated and 16 downregulated genes in females (FDR<0.05 and log2FC>0.5 or log2FC< −0.5, Fig. S3B). Sex differences were more pronounced: 82 upregulated and 40 downregulated genes (FDR<0.05 and log2FC>0.5 or FDR<0.05, log2FC< −0.5) distinguished CTL male from CTL female mice (Fig. S3C-D). As expected, upregulated genes in males included those residing on the Y chromosome (e.g. *Ddx3y*, *Uty*) and transcription factors such as *Prox2* and *Lyl1*, while downregulated genes included those located on the X chromosome (e.g. *Ddx3x*) and prolactin (*Prl*) (Fig. S3E). The effect of rearing on gene expression was sex dependent: assessing interaction among sex and rearing identified 87 genes with sex-specific regulation by rearing (Fig. 2H). GO term analysis suggested that these genes are mainly relevant to developmental processes (Fig. 2I).

### Gene translation in response to reward cues is highly rearing-dependent, modulated by sex

Because salient behavioral differences between CTL and ELA mice emerge upon exposure to reward cues (Fig. 1), we assessed reward-induced gene translation across rearing and sex in TRAPed-PVT cells (Fig. 3A). This elicited major group differences. In CTL mice, most differentially expressed genes (DEGs) were downregulated in both males and females (lists of top DEGs in S4A): 56 genes were downregulated in reward-exposed compared to no-reward males (FDR<0.05, log2FC< −0.5, Fig. 3B), and 89 genes were downregulated in reward vs no-reward females (FDR<0.05, log2FC< −0.5, Fig. 3C). In contrast, in ELA mice, most DEGs were upregulated by reward cues: 119 genes in males (FDR<0.05, log2FC>0.5, Fig. 3D), and 147 in females (Fig. 3E). Overall, exposure to reward cues changed the expression of a larger gene-set in ELA mice (Fig. 3B-E), suggesting an augmented reward response (either pro- or anhedonic) in this group.

**Fig. 3.**
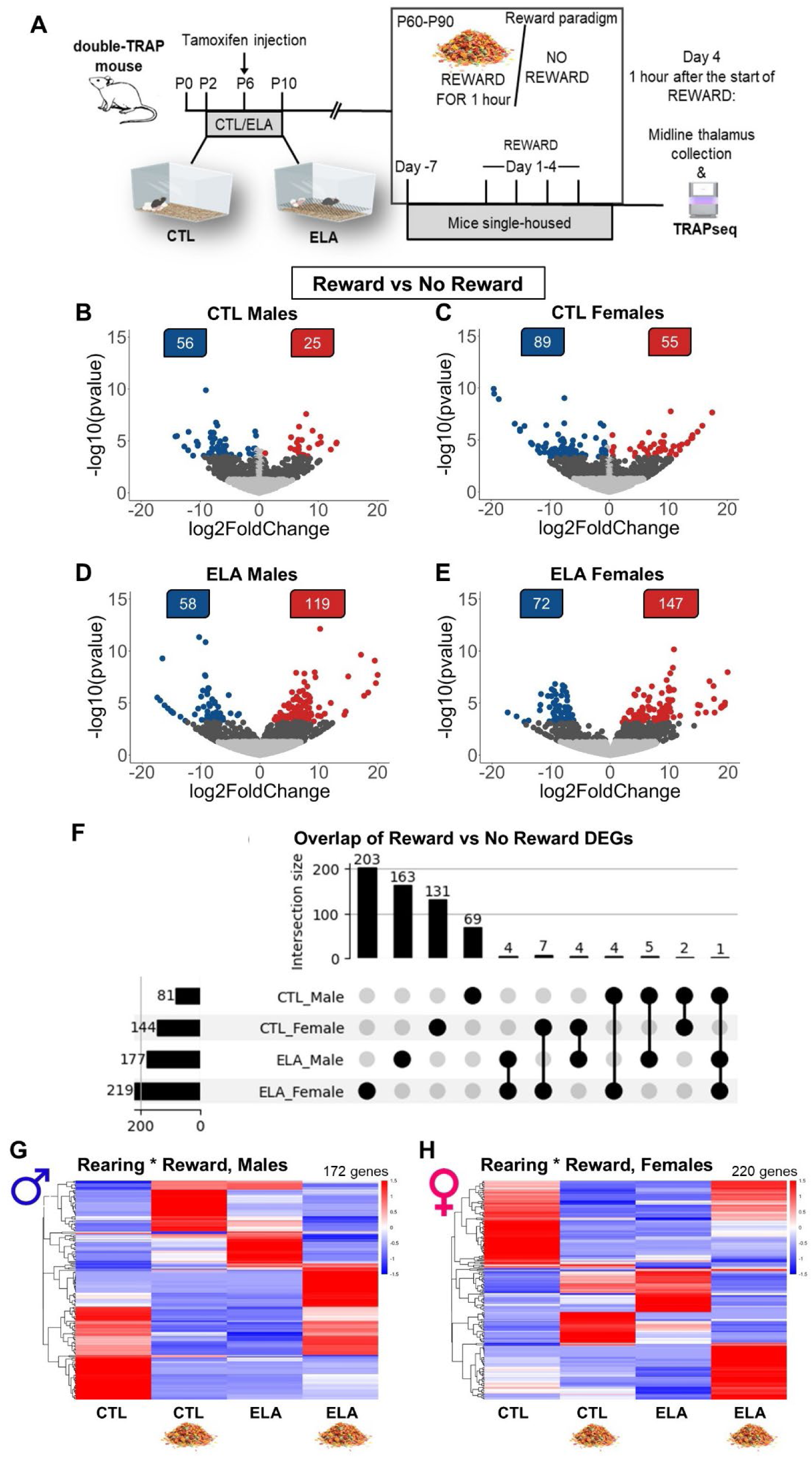
Gene translation in response to reward cues is highly rearing-dependent, modulated by sex. **A.** Experimental design for testing reward-induced changes in active translation of TRAPed-PVT cells. **B-E.** Volcano plots showing the DEGs between mice exposed to cocoa pebbles for 1 hour and mice that were not given a reward. Reward vs No Reward in CTL males (**B**), CTL females (**C**), ELA males (**D**) and ELA females (**E**). Upregulated genes shown in dark red: FDR<0.05, log2FC>0.5, downregulated genes showed in dark blue FDR<0.05, log2FC< −0.5. N = 6 No-Reward & 5 Reward CTL males, 6 No-Reward & 6 Reward ELA males, 6 No-Reward & 7 Reward CTL females, 5 No-Reward & 7 Reward ELA females **F.** Upset plot showing the number of Reward vs No Reward DEGs shared between the Rearing and Sex groups. **G-H.** Heatmaps showing averaged expression of DEGs for the interaction Rearing*Reward (FDR<0.05, log2FC>0.5 or log2FC< −0.5) in male (**G**) and female mice (**H**).

We then investigated genes and gene sets distinguishing ELA from CTL groups in their reward-induced expression, suggesting that they may contribute to the differential rearing and sex-dependent reward behaviors. Broadly, testing for the interaction of Rearing*Reward uncovered 172 genes that responded to reward cues in a rearing-dependent manner in males and 220 females (Fig. 3G-H). These results suggest that reward elicits a distinct gene expression response governed by rearing and sex, with minimal commonalities among groups.

Several genes with established brain functions were regulated by reward in more than one group. These included the potassium channel *Kcnj8* and the histone methyltransferase *Prdm9* in CTL and ELA females and the transcription regulator *Prdm15* in CTL and ELA males. In addition, the voltage-gated calcium channel *Cacna1h* was regulated in both CTL females and ELA males and the receptor tyrosine kinase *Epha3* and the neuroamidase *Neu4* in CTL and ELA males (Fig. 3F and S4B). In addition, we found an enrichment in differentially expressed transcription factors (TFs) in ELA females, and a trend in ELA males, with some TFs (e.g. *Zim1* and *Ebf2*) differentially expressed in more than one groups (Fig. S4C-E).

To identify plausible gene targets for the enduring impact of early-life adversity/ stress on adult behaviors, we combined our data-driven approach with a priori knowledge: among the DEGs, we observed that the receptor *Crhr1* for the stress-activated peptide corticotropin releasing hormone (CRH) was uniquely altered in both CTL males and females but not in either ELA group (Fig. S4B). Because of the known role of CRH-CRHR1 signaling in stress, including early-life stress (Birnie et al., 2023; Chen & Baram, 2015; Deussing & Chen, 2018; Ivy et al., 2010), we elected to further determine the role of *Crhr1* in ELA-induced disruption of reward behaviors as described below.

### Combining data- and hypothesis-driven approaches to identify candidate target genes for mediating the consequences of ELA on reward behaviors

As mentioned above, *Crhr1,* the gene encoding the CRH receptor CRHR1 was the only gene whose expression was significantly altered in opposite directions by reward cues in CTL males and females (upregulated in males and downregulated in females) and this expression change was not observed in adult ELA mice with dysregulated reward behaviors. The CRH signaling family stood out as a potential candidate because CRH is a stress- and reward-reactive peptide, and its expression is regulated by ELA in several brain regions (Brunson et al., 2001; Ivy et al., 2010; Singh-Taylor et al., 2018). In the context of ELA, CRH signaling in the nucleus accumbens associates with dampened reward behaviors in males (Birnie et al., 2023; Bolton, Molet, et al., 2018). In the PVT, a higher proportion of TRAPed-PVT cells expresses CRHR1 in ELA mice compared to CTL (Fig. 1P; Kooiker et al., 2023) and these neurons are surrounded by CRH-expressing axons (Kooiker et al., 2023; Mendoza et al., 2026), suggesting peptide-related neurotransmission. *Crh* gene translation tended to decrease in reward-exposed CTL but not ELA males (Fig. 4A), while the receptor *Crhr1* was upregulated by reward in CTL males (FDR = 0.005, Fig. 4B) and reduced in females (FDR = 0.01; Fig. 4E). A third member of the CRH signaling family, *Crhbp*, thought to sequester the peptide away from its receptor, is highly expressed in PVT (Gao et al., 2023; Shima et al., 2023) including in TRAPed-PVT neurons (Fig. 2F and Fig. 4C, F). Together with the established role of CRH signaling in stress-related plasticity (Chen et al., 2012, 2013), these results prompted further testing of altered CRH signaling in the mechanism by which ELA disrupts adult reward behaviors. To capture coordinated gene expression programs associated with rearing, reward, or the interaction between these factors, we utilized weighted gene co-expression network analyses (WGCNA). 13 distinct co-expression modules were identified in males; 2 modules were significant for rearing-specific differences and 8 modules for the interactions between rearing and reward (Fig. S5A). 20 co-expression modules emerged in females, 3 with rearing-specific differences, one with reward-specific differences and 9 with interactions between rearing and reward (Fig. S5B). *Crh*, *Crhr1*, and *Crhbp* each occupied a distinct co-expression module in both males and females. In males, the darkred module containing *Crh* was upregulated in the CTL-no-reward mice and the palegoldrenrod module containing *Crhbp* was upregulated in ELA mice compared to CTL (Fig. 4G). In females, the wheat module containing *Crhr1* was strongly upregulated in CTL no-reward mice, consistent with the rearing-dependent regulation of *Crhr1* identified by our DGEA (Fig. 4H).

**Fig. 4.**
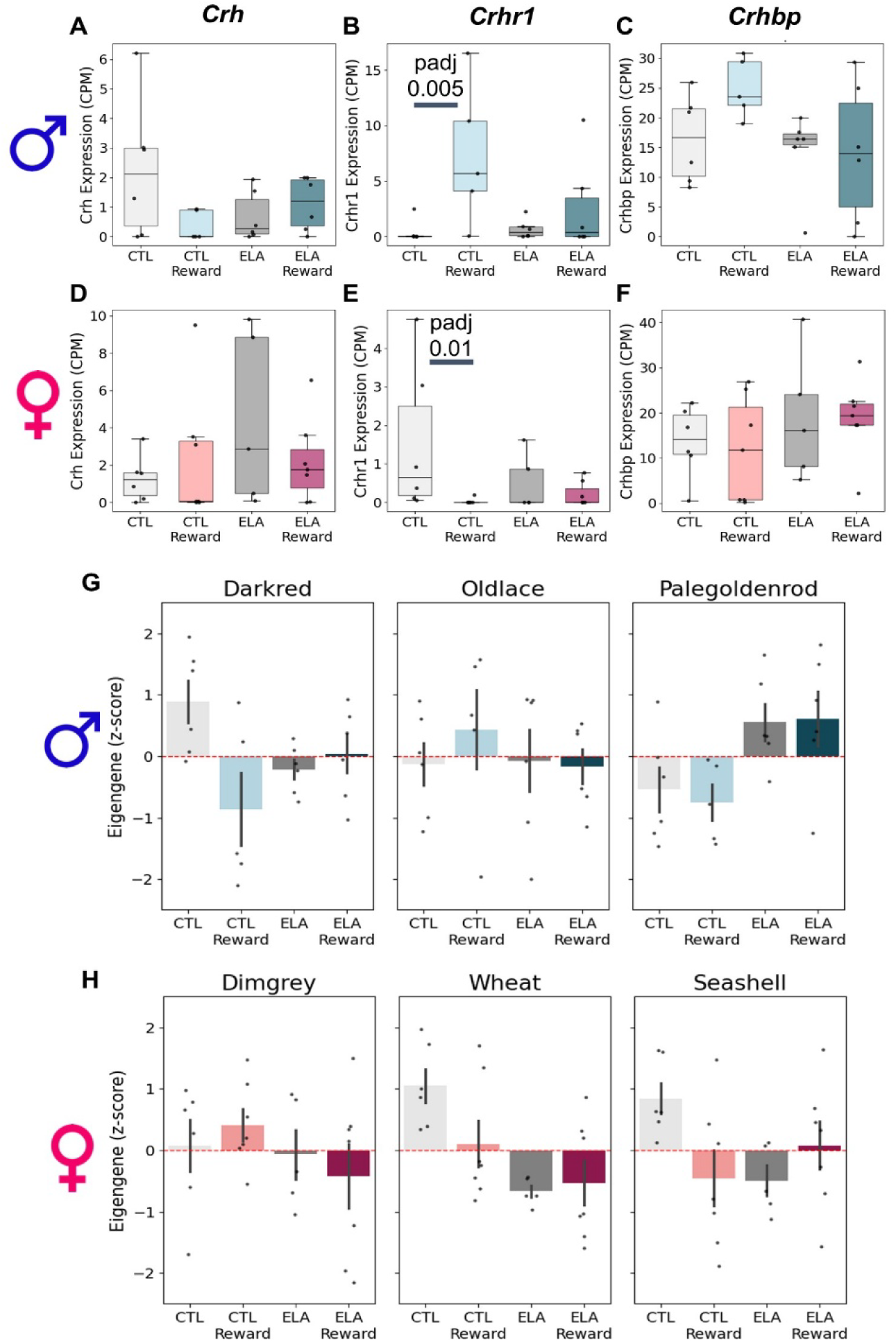
Differential gene expression and weighted gene co-expression network analyses identify CRH family genes as highly responsive to reward in a sex-dependent way, and influenced by rearing. **A-F.** Expression levels of *Crh, Crhr1* and *Crhbp* in male (**A-C**) and female mice (**D-F**) and statistical results from the DGEA. **G-H.** Module eigengene z-scores for male modules containing members of the CRH gene family plotted by condition (dark red, Crh module (5,332 genes); old lace, Crhr1 module (116 genes); pale goldenrod, Crhbp module (957 genes)) (**G**). Module eigengene z-scores for female modules containing members of the CRH gene family (dim grey, Crh module (6,100 genes); wheat, Crhr1 module (52 genes); seashell, Crhbp module (3,255 genes)) (**H**).

### *Crhr1* deletion in TRAPed-PVT cells of adult ELA mice normalizes reward behavior

Together, the above findings provided support for the hypothesis that CRH-CRHR1 signaling in TRAPed-PVT cells may contribute to the effects of ELA on reward behaviors. To test it we deleted *Crhr1* specifically in TRAPed-PVT cells of male and female CTL and ELA mice and used a within-subject design to measure reward behaviors before and after the deletion. We deleted *Crhr1* by delivering a Flp-dependent CRISPR AAV (AAV.DJ/8-CMV-Crhr1-RNA1-flexfrt-Cas9(HA)) that expresses both the CAS9 protein and a *Crhr1*-RNA-guide to the PVT of mice expressing FLPO specifically in TRAPed-PVT cells. This was accomplished by crossing TRAP2 mice with LSL-Flpo mice (B6;129S4-Gt(ROSA)26Sor^tm5(CAG-flpo)Zjh/J^, He et al., 2016) that express FLPO after CRE recombination, and administering tamoxifen to the TRAP2-Flpo offspring at P6 (Fig. 5A). We tested for *Crhr1* deletion by this intervention by quantifying CRHR1 in cells expressing the Cas9 tag HA, using single-cell resolution immunohistochemistry (Fig. 5B-C). Leveraging prior information (Kooiker et al., 2024) that inhibition of TRAPed cells in anterior PVT ameliorated ELA-induced ‘anhedonia’ in males, while inhibition of TRAPed cells in posterior PVT normalized reward behaviors in ELA females, we targeted the anterior PVT for *Crhr1* deletion in males and the posterior PVT in females.

**Fig. 5.**
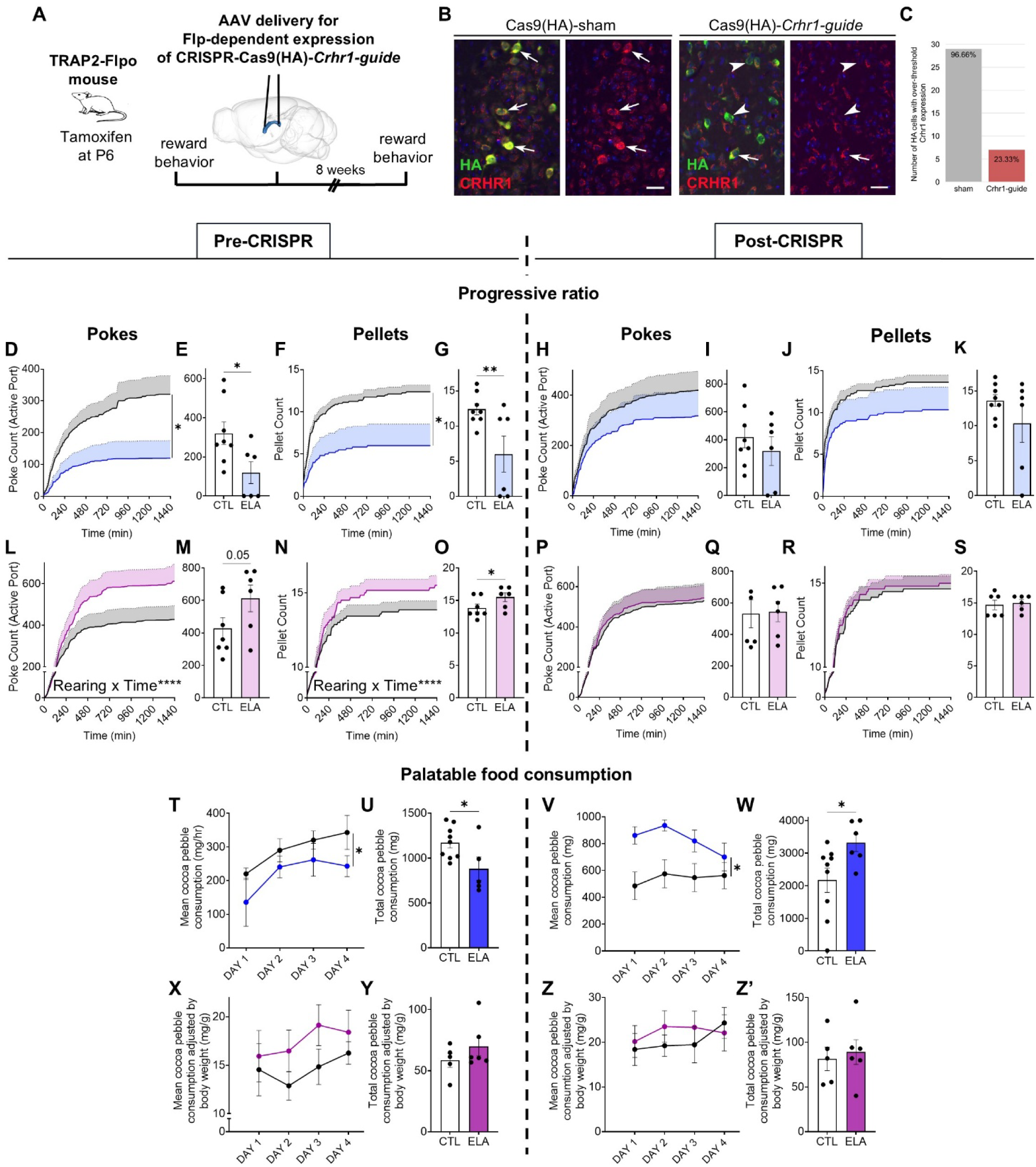
*Crhr1* deletion in TRAPed-PVT cells of adult ELA mice normalizes reward behavior. **A.** Experimental design for testing adult reward behavior before and after *Crhr1* deletion in TRAPed-PVT cells, in TRAP2-Flpo mice (*Fos-2A-iCreER^T2^* x *LSL-Flpo*). **B.** Validation of CRISPR-mediated *Crhr1* deletion in TRAPed-PVT cells. Representative microscopy images showing colocalization (or lack thereof) of CAS9-HA with CRHR1 in the PVT of mice injected with a sham virus (without *Crhr1-RNA* guides, left) or with the virus expressing *Crhr1-RNA* guide (right). Scale bars = 30 μm. Arrows show HA-CRHR1 colocalization, arrowheads show HA expression only. **C.** Quantification of cells expressing both HA and CRHR1. Cells were considered to express both HA and CRHR1 if the surface area of CRHR1 overlapped with at least 25% of the surface area of HA. **D-G.** Before Crhr1 deletion, ELA males poked the active port fewer times and received fewer sugar pellets compared to CTL, on a progressive ratio test with FED3 devices (nose pokes over 24 hours, main effect of Rearing: F(1,12) = 5.437, P = 0.0380; interaction Rearing*Time: F(1.442, 17.30) = 4.345, P = 0.0401, two-way ANOVA (**D**) and total nose pokes: t(12) = 2.426, P = 0.0160, one-tailed unpaired t test (**E**); sugar pellets retrieved over 24 hours, main effect of Rearing: F(1,12) = 3.620, P = 0.0272; interaction Rearing*Time: F(1.993, 23.92) = 4.520, P = 0.0217, two-way ANOVA (**F**) and total pellets retrieved: t(12) = 2.685, P = 0.0099, one-tailed unpaired t test (**G**)). N = 8 CTL & 6 ELA males **H-K.** After Crhr1 deletion there was no difference in number of pokes or pellets between ELA and CTL male mice (nose pokes over 24 hours, main effect of Rearing: F(1,12) = 5.863, P = 0.4586, two-way ANOVA (**H**) and total nose pokes: t(12) = 0.8024, P = 0.2190, one-tailed unpaired t test (**I**); sugar pellets retrieved over 24 hours, main effect of Rearing: F(1,12) = 1.691, P = 0.2178, two-way ANOVA (**J**) and total pellets retrieved: t(12) = 1.306, P = 0.1080, one-tailed unpaired t test (**K**)). N = 8 CTL & 6 ELA males **L-O.** Before Crhr1 deletion, ELA females poked the active port more times as the effort requirement increased, and received more sugar pellets compared to CTL, on a progressive ratio test with FED3 devices (nose pokes over 24 hours, interaction Rearing*Time: F(144, 1584) = 4.085, P < 0.0001, two-way ANOVA (**L**) and total nose pokes: t(11) = 1.774, P = 0.0518, one-tailed unpaired t test (**M**); sugar pellets retrieved over 24 hours, interaction Rearing*Time: F(144, 1584) = 2.314, P < 0.0001, two-way ANOVA (**N**) and total pellets retrieved: t(11) = 1.839, P = 0.0465, one-tailed unpaired t test (**O**)). N = 7 CTL & 6 ELA females **P-S.** After Crhr1 deletion there was no difference in number of pokes or pellets between ELA and CTL female mice (nose pokes over 24 hours, interaction Rearing*Time: F (1.295, 12.95) = 0.08776, P < 0.8335, two-way ANOVA (**P**) and total nose pokes: t(10) = 0.1228, P = 0.4524, one-tailed unpaired t test (**R**); sugar pellets retrieved over 24 hours, interaction Rearing*Time: F (2.981, 29.81) = 0.2284, P = 0.8747, two-way ANOVA (**S**) and total pellets retrieved: t(10) = 0.3627, P = 0.3622, one-tailed unpaired t test (**T**)). N = 6 CTL & 6 ELA females **T-U.** Before Crhr1 deletion, ELA males consumed less palatable food compared to CTL (cocoa pebble weight consumed/hour over 4 days, main effect of Rearing: F (1, 12) = 5.447, P = 0.0378, two-way ANOVA (**T**) and total consumption: t(12) = 2.334, P = 0.0189, one-tailed unpaired t test (**U**)). N = 9 CTL & 5 ELA males **V-W.** After Crhr1 deletion, ELA males consumed more palatable food than CTL (cocoa pebble weight consumed/hour over 4 days, main effect of Rearing: F (1, 13) = 5.128, P = 0.0413, two-way ANOVA (**V**) and total consumption: t(13) = 2.264, P = 0.0413, unpaired t test (**W**)). N = 9 CTL & 6 ELA males **X-Y.** Consumption of palatable food for CTL and ELA females before Crhr1 deletion (cocoa pebble weight adjusted by body weight, consumed/hour over 4 days (**X**) and total consumption adjusted by body weight (**Y**)). N = 5 CTL & 6 ELA females **Z-Z’.** Consumption of palatable food for CTL and ELA females after Crhr1 deletion (cocoa pebble weight adjusted by body weight, consumed/hour over 4 days (**Z**) and total consumption adjusted by body weight (**Z’**)). N = 5 CTL & 6 ELA females

Prior to CRISPR manipulations, all mice learnt to poke on the active port of FED3 devices to receive a sugar pellet (Fig. S6). In the progressive ratio test, ELA males exerted less effort and received fewer pellets compared to CTL (Fig. 5D-G), while ELA females exerted more effort and received more pellets compared to CTL (Fig. 5L-O), validating our prior cohorts (Kooiker et al., 2024; Levis et al., 2021). Following CRISPR-mediated *Crhr1* deletion, ELA-induced sex-modulated disruptions of reward behaviors were reversed, in both high effort (Fig. 5H-K and P-S) and low effort (Fig. 5T-W and 5X-Z’) tasks. Specifically, the same ELA males with low reward motivation and consumption now performed as well as CTL mice; in female, the aberrantly augmented motivation for and consumption of palatable food decreased, reaching CTL levels. Notably, behaviors of mice provided with a sham virus lacking the *Crhr1* guide RNA did not change (Fig. S8), excluding behavioral changes related to repeated testing or age, and consumption of laboratory chow did not differ between CTL and ELA mice and remained unaffected by *Crhr1* deletion (Fig. S7). Together, these results indicate that CRHR1 expressed in PVT cells activated early in life is required for the sex-dependent, ELA-induced disruption of adult reward behaviors.

## Discussion

The current studies discover key molecular mechanisms in an understudied brain region by which transient ELA leads to enduring disruptions of adult reward behaviors in a sex-specific manner. Leveraging our prior demonstration that neuronal activation within the PVT uniquely discriminates ELA from CTL rearing (Kooiker et al., 2023) and contributes to ELA-induced alterations of adult reward behavior (Kooiker et al., 2024), we 1) determine the TRAPed-PVT cell types within the heterogenous PVT, 2) identify ELA-induced enduring gene expression changes in the TRAPed cells, 3) uncover major reward-induced gene translational programs that are dictated by rearing and sex and are associated with distinct reward behaviors, 4) demonstrate that signaling via the CRH receptor CRHR1 is required for ELA-induced dysregulation of motivated adult reward behaviors in both male and female mice.

### What PVT cell types are differentially activated by experiences in the neonatal mouse?

While rearing in resource-scarce cages during postnatal days 2-9 drastically influenced reward behaviors in adult mice, how this transient exposure is converted into life-long behavioral changes is unknown. The use of genetic tagging identified quantitative differences in the number of active, FOS-expressing neurons in PVT of ELA vs CTL mice - the only regions with this discrimination among all brain regions studied (Kooiker et al., 2023). In addition, chemogenetic silencing of genetically tagged (TRAPed) cells reversed the disrupted reward behaviors of adult ELA mice (Kooiker et al., 2024), indicating that functional changes in these TRAPed-PVT cells contribute to disrupted adult reward behaviors. However, the identity of these cells remained unclear. The PVT is an elongated and highly heterogenous nucleus. Mouse PVT, unlike the human (Nishioka et al., 2026; Schulmann et al., 2024), consists exclusively of glutamatergic neurons (Frassoni et al., 1997; Kirouac, 2015; Shima et al., 2023) and the majority express calretinin, (CALB2, Mátyás et al., 2018). However, this brain node is composed of numerous neuronal cell types. These are typically clustered by expression of unique ‘marker gene’ sets into five major groups (Gao et al., 2023; Shima et al., 2023). Our data, based on bulk sequencing, suggest that TRAPed-PVT cells comprise more than one PVT-neuronal type. The use of single-cell resolution approaches indicates that the majority of TRAPed cells express calretinin (Fig. 1N) as expected, and many express NTRK1 (Fig. 1O). This places these cells within the “PVT3” cluster of Gao et al. and the “aPVT^Ntrk1^” of Shima et al. Bulk analyses of the TRAPed cells demonstrated high expression of *Htr1d, Drd1* and *Crhbp,* all markers of “PVT3” in Gao’s clustering and of *Tesc,* mainly found in the anterior-lateral PVT cluster described by Shima et al. (Fig. S2). Together, these analyses suggest that the population of cells activated early in life represents more than one specific cell type.

### In the absence of reward cues, actively translated genes in TRAPed neurons are governed mainly by sex

Actively translated (ribosome attached) genes in TRAPed-PVT cells of adult mouse PVT were minimally influenced by rearing, and varied by sex (Fig. 2H-I and S3) This was not surprising, as behaviors of mice in their home cages did not differ in ELA vs CTL mice including similar consumption of chow (Fig. S1, S7) and lack of anxiety-like behaviors (Birnie et al., 2023; Bolton et al., 2022; Taniguchi et al., 2026). Because the major behavioral differences across groups emerged upon exposure to reward or reward cues, we tested for reward-related changes in the rapid activation of gene translation across the experimental groups.

### Rewards promote major rearing and sex-dependent gene-translation programs in ELA mice of both sexes

Presentation of a palatable food reward led to changes in the translation of a large number of genes, which differed strikingly in ELA and CTL mice and in males vs females. Broadly, more genes were activated in the ELA groups (five-fold in males and three-fold in females, Fig. 3B-E). In addition, reward cues drove largely opposing changes in gene expression in CTL and ELA mice of the same sex, in accord with their distinct reward behaviors. This striking reward-induced gene translation differences of ELA vs CTL mice, in the face of modest ‘baseline’ differences suggests a latent epigenomic mechanism that is unmasked by reward presentation. Specifically, altered three-dimensional structure of the chromatin, and/ or the presence of permissive vs repressive posttranslational histone markers at promoters of salient genes might be at play, rendering the chromatin poised for reward-induced gene expression (Hokenson et al., 2026). These possibilities will be pursued in future studies.

While the abundance of actively translated genes upon reward presentation was consistent with the distinct rearing- and sex-modulated reward behaviors, they left open the question of which genes, gene-set or signaling cascades mediate the effects of ELA on PVT cell function and reward behaviors.

### The CRH receptor CRHR1 is required for ELA-induced dysregulation of motivated adult reward behaviors in both male and female mice

Aiming to identify plausible gene targets for executing the effects of ELA on adult reward behaviors, we combined the RNA sequencing data with a priori information on the neurobiology of ELA. Datawise, both the DGEA and the WGCNA identified CRH-related genes as highly responsive to reward, often changing in opposite directions between the experimental groups - for example *Crhr1* was upregulated in response to reward in CTL males and downregulated in CTL females (Fig. 4B, E). In addition, CRHR1-expressing cells were enriched in the TRAPed population in ELA vs CTL mice of both sexes. Notably, a robust body of work has implicated CRH-CRHR1 signaling in the consequences of ELA on adult brain functions (Birnie & Baram, 2025; Ivy et al., 2010; Wang et al., 2012). Thus, we reasoned that CRH-CRHR1 signaling might contribute to the sex-specific, long-lasting effects of ELA on PVT-dependent reward behavior and tested if deleting *Crhr1* in adult mice would rescue reward deficits in ELA mice, and this hypothesis is supported by the results (Fig. 5).

Whereas the data do not exclude the possibility that CRH signaling elsewhere in the brain (Birnie et al., 2023), as well as other mechanisms (e.g. Peña et al., 2017) contribute to the behavioral effects of ELA, they indicate that specific deletion of the receptor in cells activated early in life suffices to normalize reward behaviors. We also show that the deletion of *Crhr1* rather than injection of a control virus is required (Fig. S8).

Together, the current studies show that the PVT is a key encoder of adverse early-life experiences during a developmental period preceding the maturation of the hippocampal memory system. We show PVT cells active during ELA are poised for major transcriptional and translational activation upon exposure to reward cues. We finally demonstrate that manipulation of specific signaling pathways within these cells reverses of adult ELA-induced reward behavior problems, identifying potential targets for translation.

## Methods

### Animals

TRAP2 (Fos^tm2.1(icre/ERT2)Luo^, Jax #030323) and nuTRAP (Gt(ROSA)26Sor^tm2(CAG-NuTRAP)Evdr^, Jax #029899) breeders were obtained from Jackson Laboratories and bred in house. LSL-Flpo (B6;129S4-Gt(ROSA)26Sor^tm5(CAG-flpo)Zjh/J^, Jax #028584) breeders were generously provided by Dr. Josh Huang, bred in house and crossed with TRAP2 mice. All mice were housed in standard conditions at 72-75°F, 40-60% humidity and on a 12-h light-dark cycle (lights on at 7 AM). Mice used in experiments were 2-7 months old. 2-4 weeks prior to experiments, mice were transferred to a reverse light-dark cycle room (lights on at 12 AM). All behavioral experiments started within 2 hours of the start of the active, dark cycle. Mice were single-housed following surgical procedures or prior to assessing reward behavior. All experimental procedures were approved by the University of California-Irvine Institutional Animal Care and Use Committee (AUP 24-104, 21-128 and 18-183) and were in accordance with the guidelines from the National Institute of Health.

### Limited bedding and nesting and activity-dependent genetic labeling

Pregnant dams were single-housed on embryonic day 17 (E17) and monitored for birth of pups. Our ELA paradigm - limited bedding and nesting (LBN) - started on the morning of P2. First, litters were culled to a maximum of 6 pups including both sexes. CTL dams and pups were placed in cages with a standard amount of corn cob bedding (400 mL) and cotton nestlet material. ELA dams and pups were placed in cages with one half cotton nestlet placed on a fine-gauge plastic-coated aluminum mesh platform, 2.5-cm above the floor of the cage, covered with sparse corn cob bedding. Dams and pups were returned to standard cages on the morning of P10. Activity-dependent genetic labeling was induced with tamoxifen administration at P6. Pups were removed from the cage for a duration of less than 10 minutes, placed on a warming pad, and injected subcutaneously with 60 mg/kg tamoxifen (catalog no.T5648; MilliporeSigma) dissolved in corn oil (catalog no. C8267; MilliporeSigma).

### Behavioral testing

#### Palatable food consumption

Single-housed mice were habituated to ∼1 g of Cocoa Pebbles cereal (Post, USA) in their home cage overnight (Day 0). On days 1-4, pre-weighed cocoa pebbles (∼1g) were placed in their homecage, retrieved one hour later, and weighed again to calculate the intake. Mice were weighed on Day 0.

#### Motivational testing

Feeding Experimentation Devices (FED3) (Matikainen-Ankney et al., 2021) were programmed with a “Fixed Ratio” where the device dispenses 1 sugar pellet (TestDiet 1811149, sucrose tab/chocolate 20mg) following 1 nose-poke on the active port. The devices were then placed in the homecage of single-housed mice for 48 hours (training phase, Day 0-2). On Day 3, the FED3 devices set on the same fixed ratio were placed in the homecages for 4 hours to assess whether mice had learnt to nose-poke the active port to retrieve sugar pellets (learning test phase). Then, the FED3 devices were programmed with a progressive ratio, where mice need to poke increasingly more times to receive the same reward (1 sugar pellet). The progressive ratio schedule used the series 1, 2, 4, 6, 9, 12, 15, 20, 25, 32, 40, 50, 62, 77, 95, 118, 145, 178, 219 etc., according to the formula by Richardson and Roberts, 1996. On Day 4, the devices were placed back in the cages for 24 hours (motivational testing phase) and then the data were retrieved and analyzed. Each device was returned to the same mouse throughout the experiment. The devices were placed in the cages with 1 pellet on the tray and few (∼5) pellets in front of the tray to encourage nose-pokes.

### Translating ribosome affinity purification and RNA sequencing (TRAPseq)

#### Tissue collection

The mice were euthanized by decapitation one hour after the initiation of reward on Day 4 (reward experiment) or at a designated time of day without any prior disturbance (baseline experiment). The brain was extracted and the midline thalamus mainly composed of the whole PVT was extracted on ice, rapidly frozen on dry ice and stored in −80℃ until further processing.

#### TRAPseq

Actively translated RNA was isolated with the TRAP protocol as described in (Heiman et al., 2014). The samples were homogenized with a 2 mL glass potter tissue grinder and PTFE pestle (Duran Wheaton Kimble 358029), on a Rotor for homogenizers (Yamato LT-400D) in 500uL tissue lysis buffer. The affinity matrix was prepared using 37.5 μL of Streptavidin MyOne T1 Dynabeads (Invitrogen 65601) per sample (the optimal bead titer was determined with an affinity matrix titration), with the corresponding volumes of biotinylated protein L and the GFP antibodies 19C8 and 19F7 (Memorial Sloan-Kettering Monoclonal Antibody Facility), in low-salt buffer. The affinity matrix (final volume 100 μL/sample) was added to every sample for immunopurification (IP), and samples were incubated on a tube rotator at 4℃ for roughly 18 hours. For the pre-IP samples, a small fraction (5% of the total volume) was aliquoted from every sample before the IP start (i.e. before the addition of the affinity matrix) and stored at 4℃ for the duration of the IP. After IP and washes in high-salt buffer, the RNA from both post-and pre-IP samples was eluted and purified with the RNeasy Plus Micro Kit (74034, Qiagen). The RNA was first dissociated from the ribosomes/beads with 350 μl RLT Plus containing 40 μM dithiothreitol (DTT, Thermo Fisher Scientific R0861), and purified according to the manufacturer’s instructions. The Smart-seq2 protocol (Picelli et al., 2014) was used for cDNA conversion, amplification and library preparation with Nextera XT DNA Library Preparation Kit (Illumina FC-131-1096). The RNA and DNA concentrations were measured with the Qubit™ 1X RNA High Sensitivity (HS) Broad Range (BR) and Extended Range (XR) Assay Kit, 500 assays (Thermo Fisher Scientific Q32855) and the dsDNA High Sensitivity (HS) and Broad Range (BR) Assay (Thermo Fisher Scientific Q33231) respectively, on a Qubit™ 4 Fluorometer, with WiFi (Thermo Fisher Scientific Q33238). The RNA integrity and library profiles were evaluated with the Agilent RNA 6000 Pico Kit (Agilent 5067-1513) and High Sensitivity DNA Kit (Agilent 5067-4626) respectively, on a 2100 Bioanalyzer (Agilent Technologies Inc., CA). The libraries were sequenced on a NextSeq2000 platform as 100 bp paired-end reads to at least 20 M raw read depth.

#### Data analysis

Adapters (CTGTCTCTTATACACATCT) were trimmed using cutadapt (Martin, 2011) and reads were aligned with STAR (Dobin et al., 2013) using the mm10 genome with Gencode vM21 annotations. Differential gene expression analysis (DGEA) was performed with a negative binomial generalized linear model and the Benjamin-Hochberg correction to calculate false discovery rates (FDR) (DESeq2, Love et al., 2014), after correction for potential batch and litter effects with ComBat-seq (Zhang et al., 2020). For the post- vs pre-IP comparison with DESeq2, genes were retained if they had expression levels of at least 10 counts in 10 samples. For all other comparisons (Rearing, Sex, Reward) genes were retained if they had expression levels of at least 10 counts in 3 samples.

Weighted gene co-expression network analysis (WGCNA) was performed using PyWGCNA rezaie(Rezaie et al., 2023). Analyses were conducted separately for males and females using ELA and CTL, and Reward and No-Reward samples. Only protein-coding genes were included, and genes were retained if they had expression levels of at least 1 TPM in a minimum of three samples. The same network construction parameters were applied to both sexes, including a minimum module size of 50 and a MEDissThres of 0.6. Module-trait correlations were then computed to identify modules significantly associated with condition. Eigengene values were calculated by averaging normalized expression across all genes in a module.

### Virus delivery in the PVT

The viruses used for the CRISPR-mediated *Crhr1* deletion (AAV.DJ/8-CMV-crfr1-RNA1-flexfrt-Cas9-HA, titer 1.92E+13 gc/mL, and AAV.DJ/8-CMV-crfr1-RNA2-flexfrt-Cas9-HA, titer 1.56E+13 gc/mL, mixed 1:1) and sham experiments (AAV.DJ/8-CMV-empty-flexfrt-Cas9-HA, titer 1.85E+13 gc/mL) were designed by Dr. Nicholas Justice and produced by the Gene Vector Core, Advanced Technology Core, Baylor College of Medicine. For virus delivery, the mice were anesthetized with isofluorane and placed in a stereotactic frame (Robot Stereotaxic instrument – StereoDrive, NEUROSTAR). The virus was aspirated in a glass capillary (Calibrated Pipets, Drummond Scientific Company 2-000-010) connected to a slow injector (PICOSPRITZER III - Intracellular Microinjection Dispense Systems, Parker 052-0500-900). 400 μL of virus was infused bilaterally in the anterior PVT (AP −0.41, ML +/−0.37, DV +3.55 at 6° angle) of male mice and in the posterior PVT of female mice (AP −1.50, ML +/−0.30, DV +2.90 at 6° angle), over 12-15 minutes. The health of the mice was monitored closely for the next 5 days. Post-CRISPR experiments took place roughly 2 months after virus delivery, to allow for virus expression and effective removal of the CRHR1 receptor.

### Immunohistochemistry and imaging

#### Tissue collection and processing

Mice were euthanized with sodium pentobarbital and perfused transcardially with saline at room temperature (RT) for 2 minutes, followed by 4% paraformaldehyde in 0.1M sodium phosphate buffer (PB, pH = 7.4) (4% PFA in PB, RT) for 15 minutes. The brains were removed and postfixed in 4% PFA for 3-6 hours, and then cryoprotected in 15%, followed by 30% sucrose solution in 0.1 M PB. Brains were frozen and cryosectioned coronally at a 20 μm or a 35 μm thickness using a cryostat (Leica Biosystems CM1900). The sections were collected in 0.1 M PB for immunohistochemistry or stored in an anti-freeze solution.

#### Immunohistochemistry

For fluorescent immunolabeling of CRHR1 in Fig. 1, free-floating sections were washed and permeabilized in PBS containing 0.3% Triton X-100 (PBS-T) for 15 minutes (3 washes of 5 minutes each). Sections were incubated in 5% normal donkey serum (NDS) for 1 hour. Next, the sections were incubated in primary antibody solution containing goat anti-CRHR1 (1:2,000, aa107-117, N-terminal, Everest Biotech, EB08035, Lot P1 E090910), at 4°C for 3 days. Subsequently the sections were washed again in PBS-T and incubated in secondary antibody solution containing Cy^TM^3-conjugated affinipure donkey anti-goat IgG (H+L) (1:400, Jackson ImmunoResearch, AB_2340411) or donkey anti-Goat Alexa Fluor 488 (1:400, Invitrogen, A-11055) in PBS-T (RT), in the dark, under agitation for 2 hours. Finally, the brain sections were washed again, mounted on gelatin-coated slides, and coverslipped with a slide mount for fluorescent labeling (Bioenno, Cat# 032019 or Cat# 032347) or Vectashield mounting medium with DAPI (Vectashield, Vector Laboratories H-1200).

The immunolabeling of CALB2 and NTRK1 was performed as described above. Sections were incubated with a mouse monoclonal anti-CALB2 (1:5,000, Chemicon, MAB1568) or a rabbit polyclonal anti-TrkA (also known as NTRK1) (1:1,000, Sigma, cat# 06-574) for 3 days (4°C), and visualized using anti-mouse or anti-rabbit IgG conjugated to Alexa Fluor 568 (1:400, Invitrogen), respectively.

For fluorescent labeling of cFos, free-floating sections were first washed in PBS-T, incubated in 5% normal goat serum (NGS) in PBS-T for 1 hour and then in primary antibody solution containing rabbit anti-cFos (1:20,000, Oncogene, Ab-5, PC38) for 3 days at 4°C. Following 3 x 5 minutes washes in PBS-T, the sections were incubated in secondary antibody solution with goat anti-rabbit Alexa Fluor 568 (1:400, Invitrogen A-11011) for 2 hours at room temperature. Sections were then washed 3 x 5 min in PBS-T, mounted on gelatin-coated slides, and coverslipped with a slide mount (Bioenno, Cat#032019).

For immunolabeling of CRHR1 in Fig. 5, we targeted the C-terminus of the protein, as the RNA guides used are specific to the DNA sequence corresponding to this site. Free-floating sections were incubated in goat anti-CRHR1 (1:5,000, Santa Cruz Biotechnology Inc., C-20, sc 1757) for 3 days (4°C), followed by washing in PBS-T (3 x 5 minutes) and incubating in secondary antibody solution containing donkey anti-goat Alexa Fluor 568 (1:400, Invitrogen, A-11057) (2 hours, RT). For dual labeling, sections were further washed 3 x 5 min in PBS-T, incubated in a rabbit polyclonal anti-HA (1:4,000, Sigma, H6908) for 2 days (4°C), and visualized using anti-rabbit Alexa Fluor 488 (1:400, Invitrogen, A21206). The sections were mounted on gelatin-coated slides and coverslipped with a slide mount (Bioenno, Cat#032019).

#### Image acquisition

Confocal images were acquired using an LSM-510 confocal microscope (Zeiss) with an apochromatic 10X, 20X, or 63X objective. Virtual z sections of 1 mm were taken at 0.2- to 0.5 mm intervals. Image frame was digitized at 12-bit using a 1024 X 1024 pixel frame size.

### Statistical analysis

During all experiments, mice from different experimental groups were tested (or the tissue was collected) in a randomized order, and the experimenters were blinded to the experimental groups. The number of biological replicates (N) is indicated in the figures (one data point is one subject) and in the figures legends, and refers to the total number of experimental subjects independently treated in each condition. Averaged data are presented as mean values, accompanied by standard error of mean (SEM). Subjects were excluded from analyses if they were statistical outliers with the Grubb’s test (Alpha = 0.05), or in case of technical problems during data acquisition.

For behavioral tests and immunohistochemistry, GraphPad Prism 10 was used for statistical analyses. We used independent samples t tests when comparing two independent groups, and paired-samples t tests for within subject comparisons. We used two-way ANOVAs for two-factorial designs (e.g. Time x Rearing). Significant main effects and interactions were analyzed using Sidak’s post hoc tests. For the RNA sequencing data, DGEA analysis was performed in Rstudio (RStudio 2025.05.0, R version 4.5.0) with the package DESeq2 (Love et al., 2014).

## Author contributions

TZB and AFS conceived the project, designed the experiments and wrote the manuscript. AFS performed experiments (stereotactic virus delivery, behavioral testing), tissue processing (tissue collection, TRAP, TRAPseq) and data analysis and interpretation. RW performed data analysis and interpretation and wrote the manuscript. YC performed tissue processing, IHC and data analysis. CLK, MTB and AKS contributed to study design. MRS and RR contributed to experiments and tissue processing. HYL and MG contributed to library preparation, RNA sequencing and data analysis. AM contributed to study design, data analysis and interpretation.

## Acknowledgements

We thank Graciella Angeles, Lia Harvey and Jennifer Daglian for assistance with animal breeding and housekeeping. We also thank Ku Kim, Lauren R. Sayat, Ali Zaidi and Thinh Bao Phan for technical support.

## Supplementary Figures

**Fig. S1.**
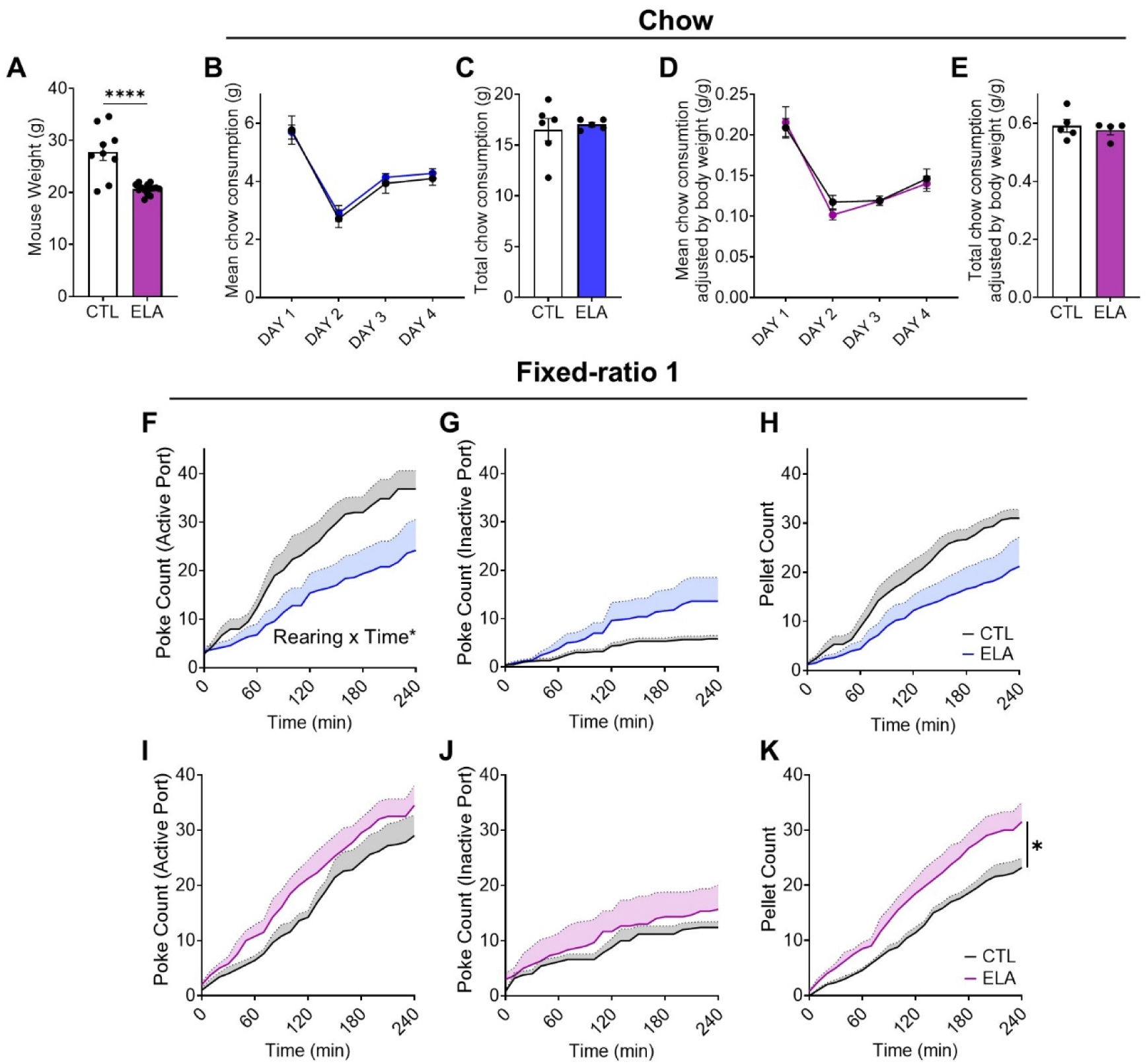
**Related to Fig. 1**. **A.** ELA female mice weighed less than CTL (t(21) = 5.344, P < 0.0001, unpaired t test). **B.** Mean daily chow consumption of CTL and ELA male mice. **C.** Total chow consumption of CTL and ELA male mice over the course of 4 days. **D.** Mean daily chow consumption of CTL and ELA female mice. **E.** Total chow consumption of CTL and ELA female mice over the course of 4 days. **F-H.** Performance of CTL and ELA male mice on a 4-hour fixed-ratio 1 test on FED3 devices. Mice poked the active port (**F**) more times than the inactive port (**G**) to receive sugar pellets (**H**) (effort = 1 nose poke on the active port for the reward = 1 sugar pellet). ELA males poked the active port fewer times compared to CTL mice over time (nose pokes over 4 hours, interaction Rearing*Time: F (2.214, 19.93) = 3.542, P = 0.0443; main effect of Rearing: F (1, 9) = 3.866, P = 0.0808, two-way ANOVA (**F**)). **I-K.** Performance of CTL and ELA female mice on a 4-hour fixed-ratio 1 test on FED3 devices. Mice poked the active port (**I**) more times than the inactive port (**J**) to receive sugar pellets (**K**) (effort = 1 nose poke on the active port for the reward = 1 sugar pellet). ELA females retrieved more sugar pellets compared to CTL (main effect of Rearing: F (1, 7) = 7.551, P = 0.0286, two-way ANOVA (**K**)).

**Fig. S2.**
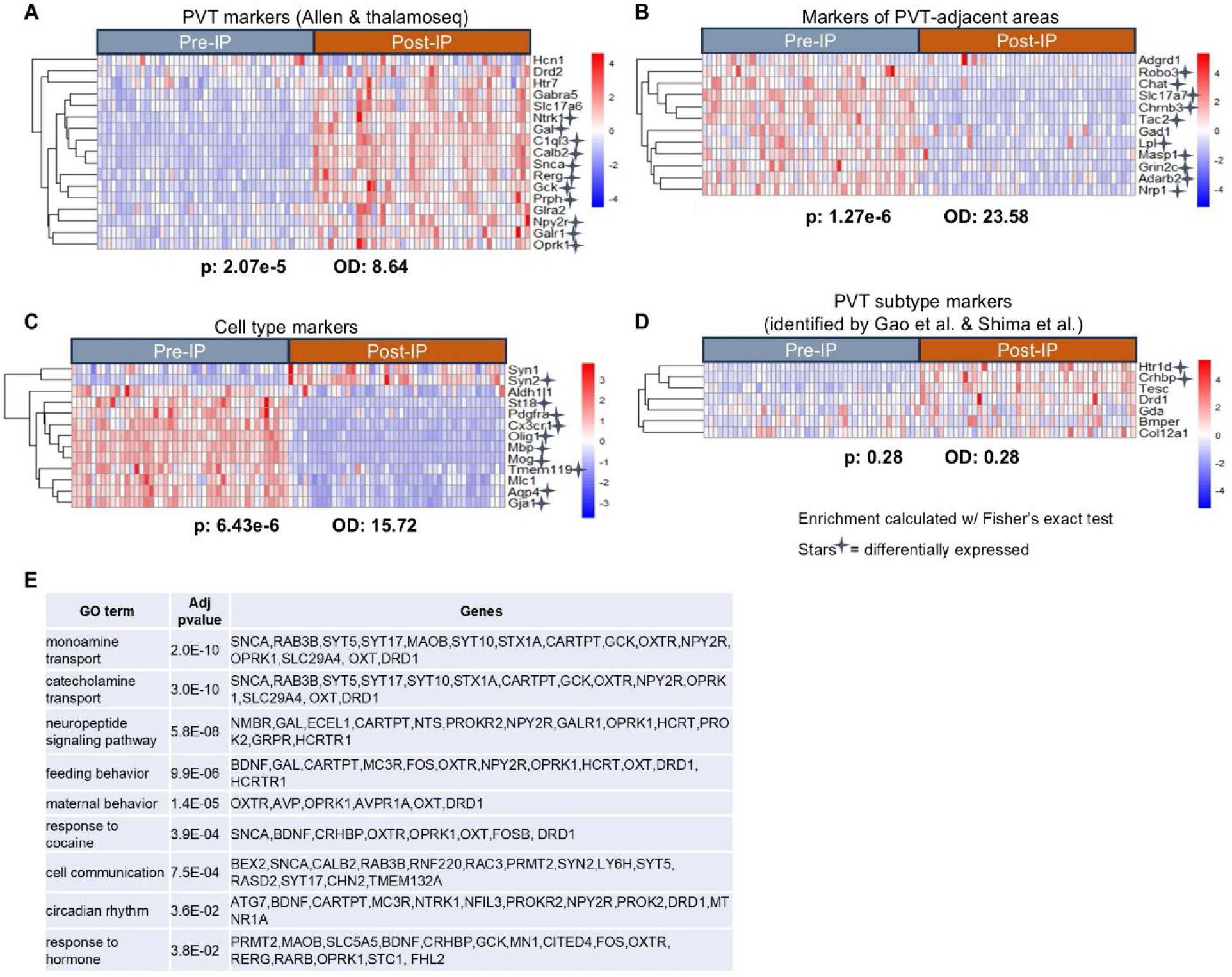
**Related to Fig. 2**. **A-D.** Heatmaps showing expression levels of cell-type or neuronal subtype markers, and statistical results. **E.** GO term analysis for the DEGs upregulated in the post-IP fraction (FDR<0.05, log2FC>1) (Fig. 2F).

**Fig. S3.**
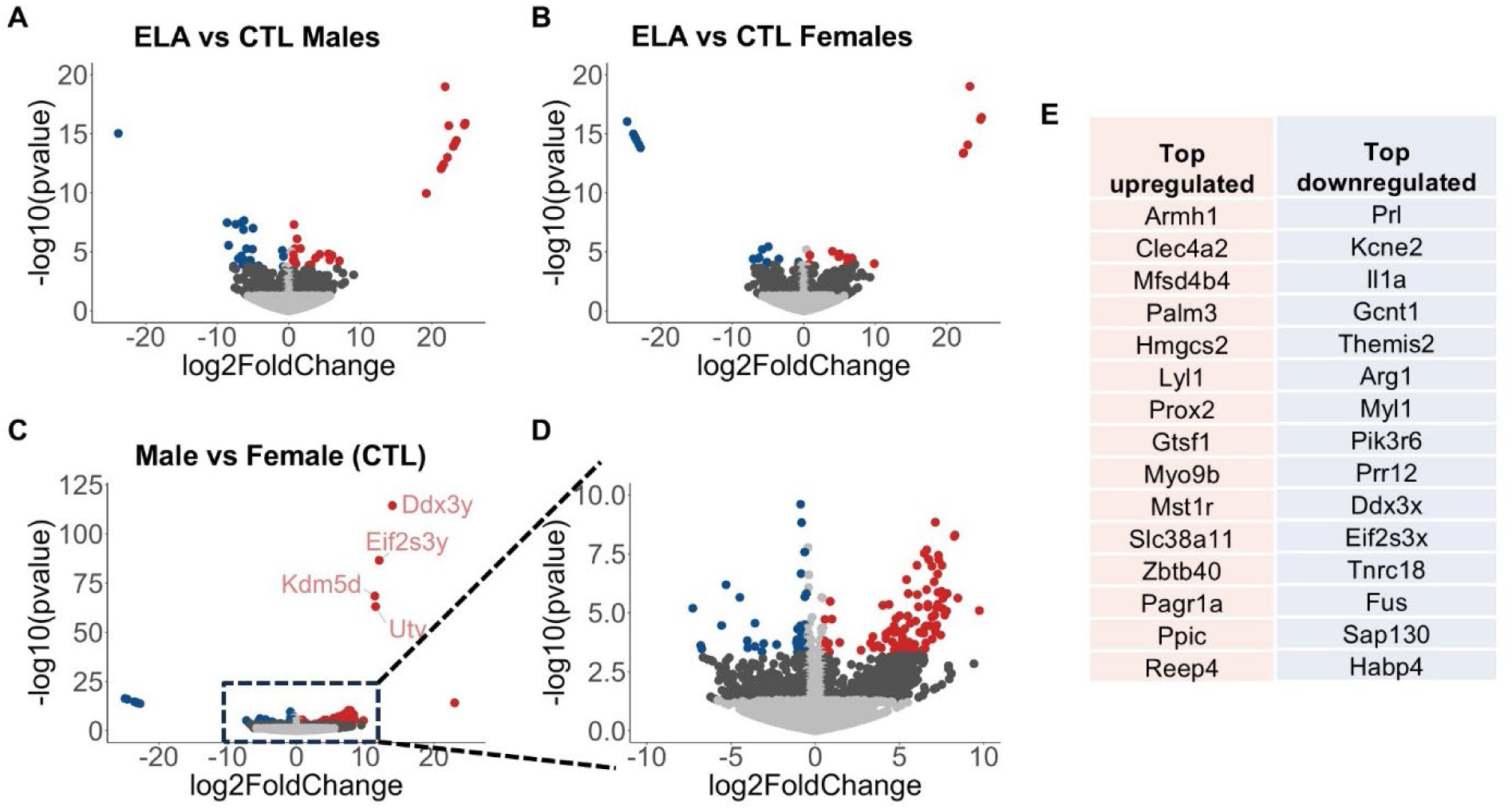
**Related to Fig. 2**. **A-B.** Volcano plots showing the DEGs between ELA and typical rearing (CTL) in male (**A**) and female (**B**) mice. **C-D.** Volcano plots showing the DEGs between CTL male and female mice. **E.** Top 15 upregulated and downregulated sex-dependent DEGs shown in the volcano plot in (**D**). Upregulated genes shown in dark red: FDR<0.05, log2FC>0.5, downregulated genes showed in dark blue FDR<0.05, log2FC< −0.5. N = 10 CTL & 13 ELA males, 12 CTL & 10 ELA females

**Fig. S4.**
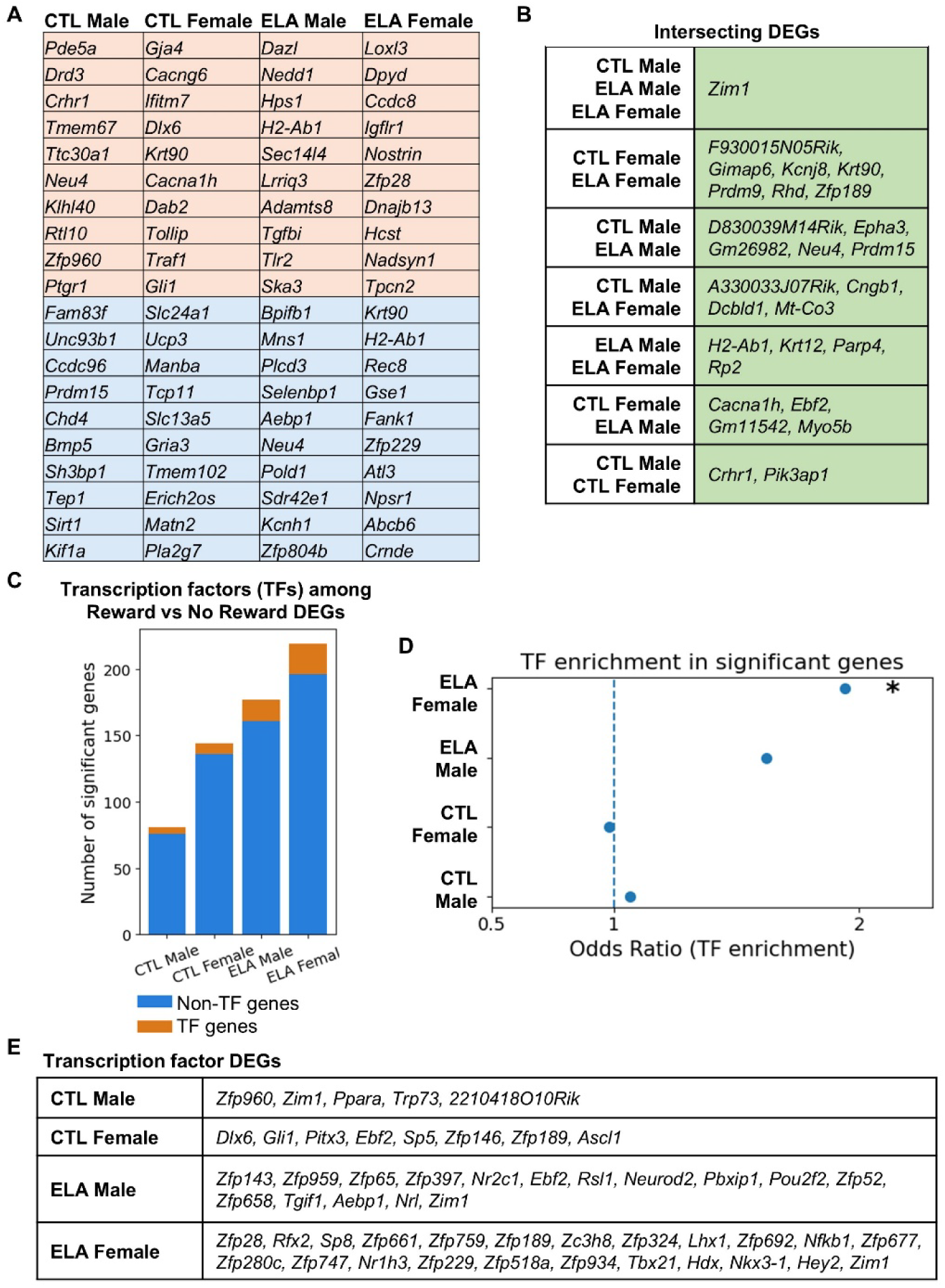
**Related to Fig. 3**. **A.** Top 10 (with the lowest FDR) up- and downregulated Reward vs No-Reward DEGs for every experimental group. **B.** Lists of DEGs that are overlapping between experimental groups, shown in the upset plot in Fig. 3F. **C.** Counts of DEGs identified in Fig. 3B–E, grouped by annotation as transcription factors (TFs) versus non–TFs. **D.** Fisher’s exact test for the enrichment of TFs among DEGs. ELA females show a ∼2x enrichment for TFs (odds ratio = 1.94, P = 0.0051) and ELA males show a trend (odds ratio = 1.62, P = 0.074). **E.** TFs with differential gene expression in Reward vs No-Reward, in every experimental group.

**Fig. S5.**
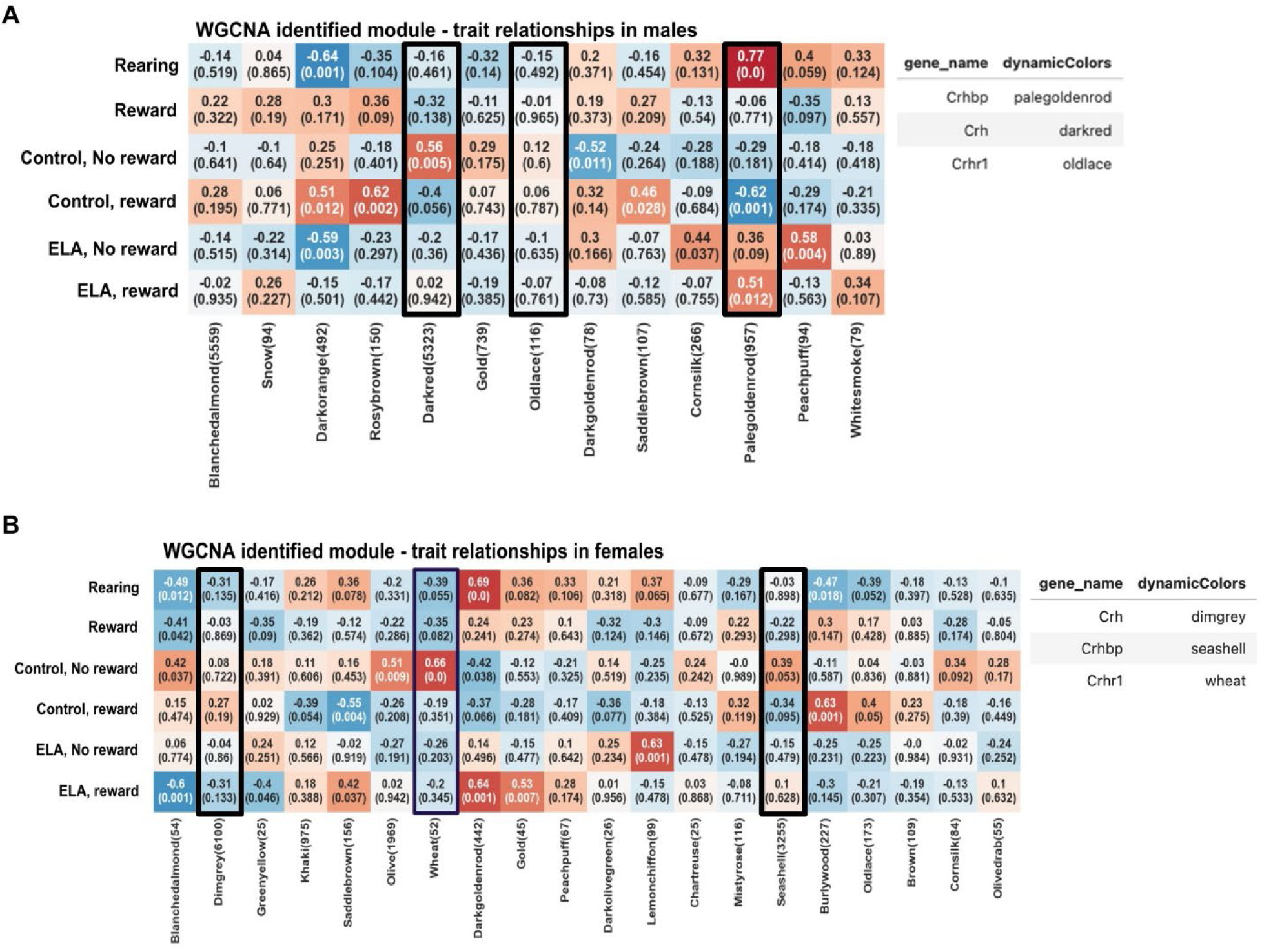
**Related to Fig. 4**. **A-B.** Module-trait relationship heatmaps generated using PyWGCNA for ELA/CTL and Reward/No-Reward conditions in males (**A**) and females (**B**). Numbers in parentheses indicate the significance (p-value) of the module-trait correlation. Color represents the correlation coefficient (red, positive correlation; blue, negative correlation). Modules containing CRH family genes are outlined in black. In males: dark red (*Crh*), old lace (*Crhr1*), and pale goldenrod (*Crhbp*). In females: dim grey (*Crh*), wheat (*Crhr1*), and seashell (*Crhbp*).

**Fig. S6.**
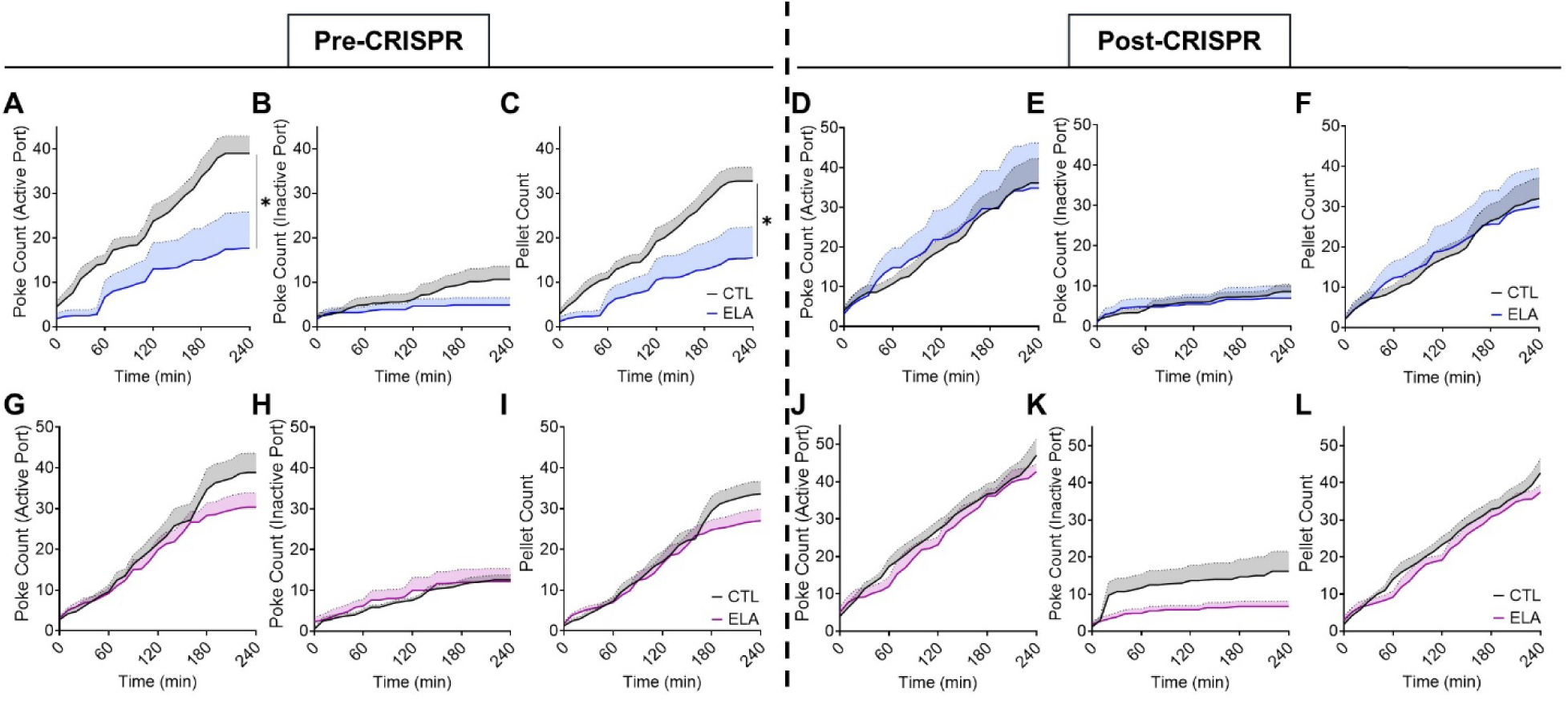
**Related to Fig. 5**. **A-F.** Performance of CTL and ELA male mice on a 4-hour fixed-ratio 1 test, on FED3 devices. Mice poked the active port (**A, D**) more times than the inactive port (**B, E**) to receive sugar pellets (**C, F**) (effort = 1 nose poke on the active port for the reward = 1 sugar pellet) both before and after *Crhr1* deletion. Before *Crhr1* deletion, ELA males poked the active port fewer times compared to CTL mice (main effect of Rearing: F(1, 12) = 6.146, P = 0.0290; interaction Rearing*Time: F(1.531, 18.37) = 3.968, P = 0.0464, two-way ANOVA (**A**)) and received fewer sugar pellets (main effect of Rearing: F(1, 12) = 5.641, P = 0.0351; interaction Rearing*Time: F(1.417, 17.00) = 4.174, P = 0.0451, two-way ANOVA (**C**)), while there was no significant difference between CTL and ELA males after *Crhr1* deletion (**D-F**). **D-F.** Performance of CTL and ELA female mice on a 4-hour fixed-ratio 1 test on FED3 devices. Mice poked the active port (**G, J**) more times than the inactive port (**H, K**) to receive sugar pellets (**I, L**) both before and after *Crhr1* deletion.

**Fig. S7.**
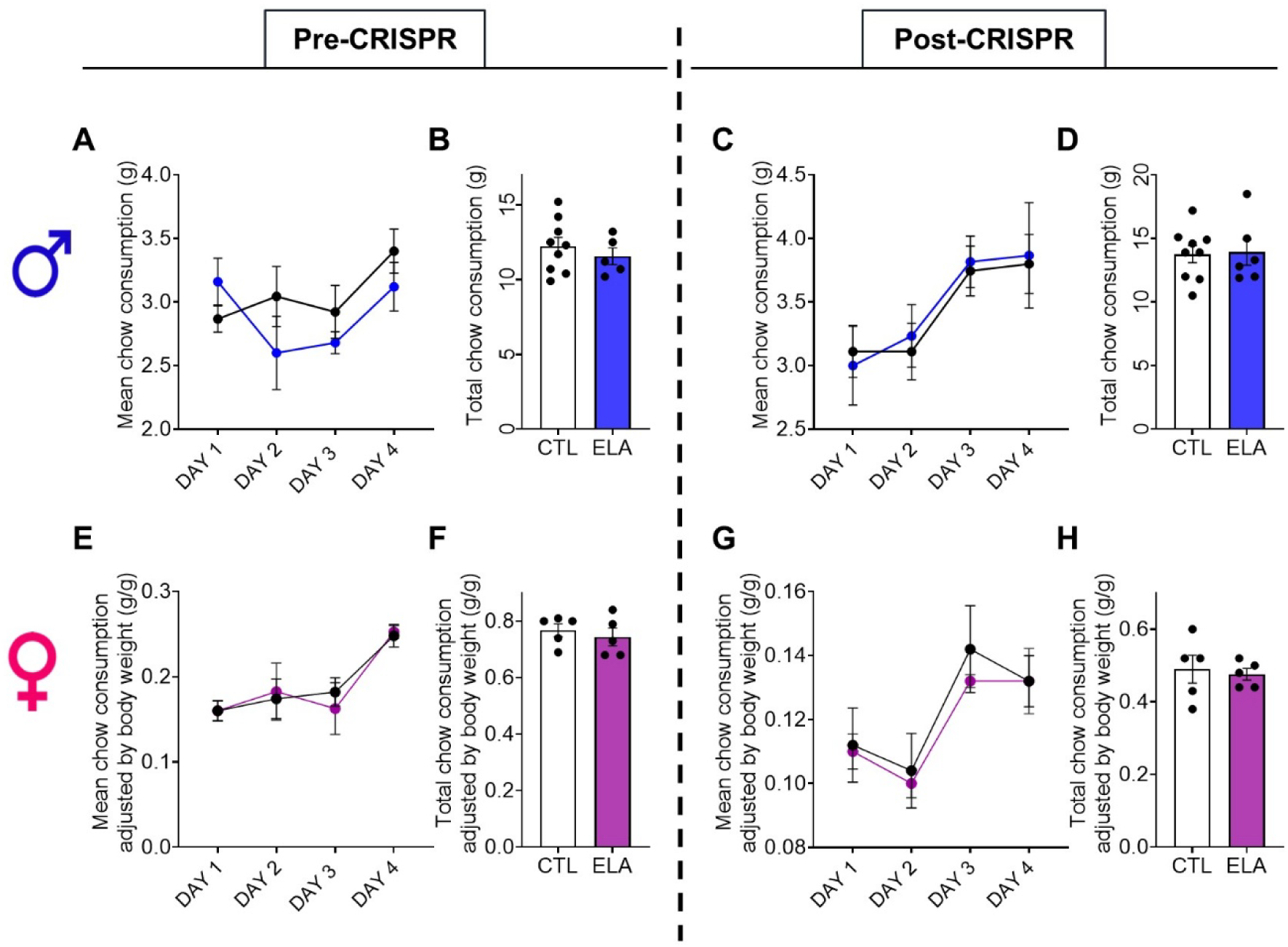
**Related to Fig. 5**. **A-D.** Chow consumption of CTL and ELA males before (**A-B**) and after (**C-D**) CRISPR-mediated *Crhr1* deletion. Daily mean consumption (**A, C**) and total consumption over 4 days (**B, D**). **E-H.** Chow consumption of CTL and ELA females before (**E-F**) and after (**G-H**) CRISPR-mediated *Crhr1* deletion. Daily mean consumption (**E, G**) and total consumption over 4 days (**F, H**).

**Fig. S8.**
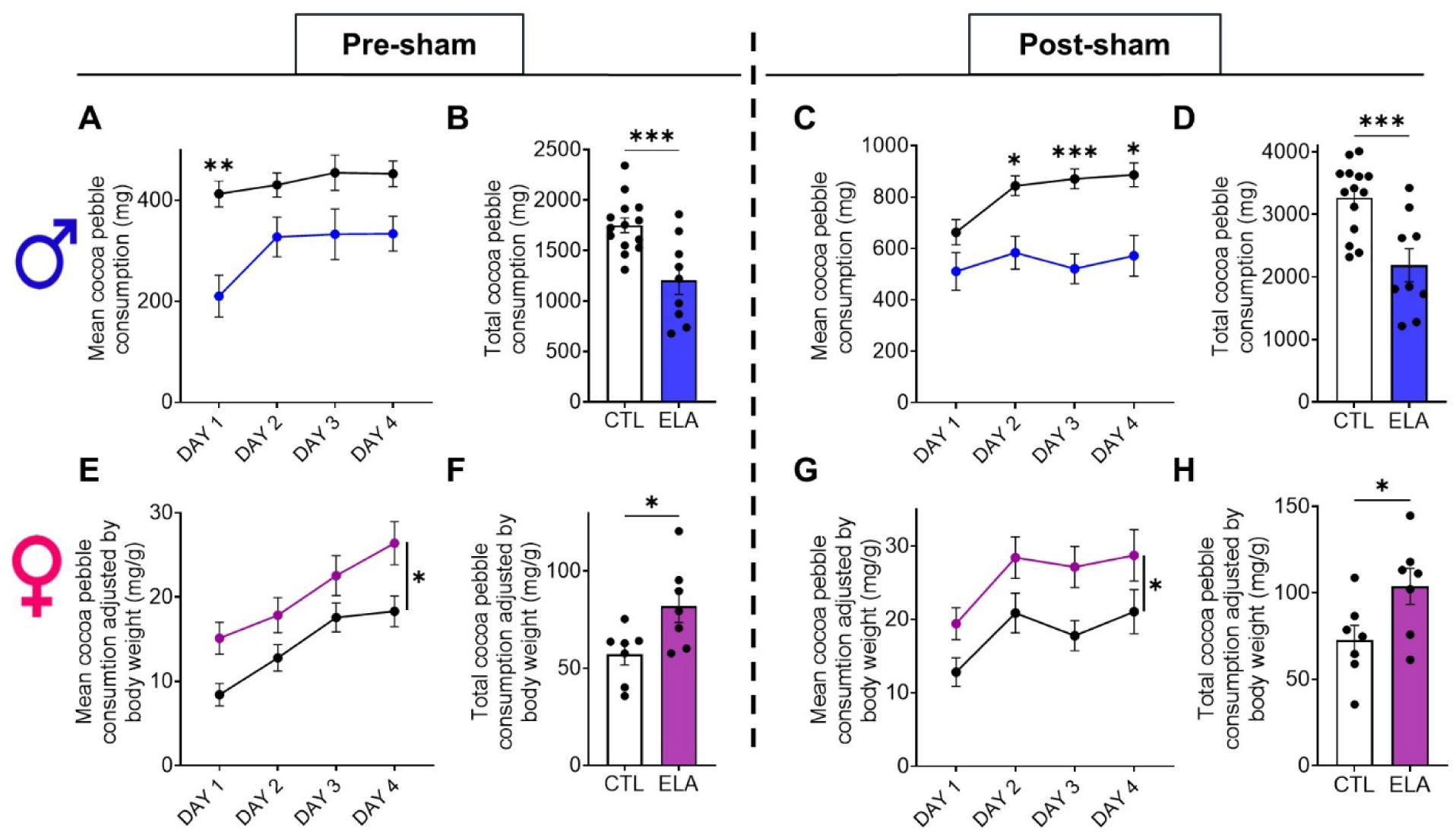
**Related to Fig. 5**. **A-B.** Before the expression of the sham virus (a virus that expresses CAS9-HA but no RNA guides), ELA mice consumed less palatable food compared to CTL mice (cocoa pebble weight consumed/hour over 4 days, main effect of Rearing: F(1, 21) = 14.59, P = 0.0010, two-way ANOVA with Sidak post hoc tests (**A**) and total consumption: t(21) = 3.820, P = 0.0010, unpaired t test (**B**)). **C-D.** After the expression of the sham virus ELA mice consumed less palatable food compared to CTL mice (cocoa pebble weight consumed/hour over 4 days, main effect of Rearing: F(1, 21) = 14.60, P = 0.0010; interaction Rearing*Day: F(2.474, 51.95) = 5.692, P = 0.0034, two-way ANOVA with Sidak post hoc tests (**C**) and total consumption: t(21) = 3.821, P = 0.0010, unpaired t test (**D**)). **E-F.** Before the expression of the sham virus, female ELA mice consumed more palatable food compared to CTL (cocoa pebble weight adjusted by body weight, consumed/hour over 4 days, main effect of Rearing: F(1, 12) = 6.273, P = 0.0277, two-way ANOVA (**E**) and total consumption adjusted by body weight: t(12) = 2.505, P = 0.0277, unpaired t test (**F**)). **G-H.** After the expression of the sham virus, female ELA mice consumed more palatable food compared to CTL (cocoa pebble weight adjusted by body weight, consumed/hour over 4 days, main effect of Rearing: F(1, 12) = 5.256, P = 0.0407, two-way ANOVA (**G**) and total consumption adjusted by body weight: t(12) = 2.293, P = 0.0407, unpaired t test (**H**)). N = 14 CTL & 9 ELA males, 7 CTL & 7 ELA females

